# hPSC-Derived Distal Lung Organoids Reveal Respiratory Airway Secretory Cells Acting as Immune Sentinels in Human Distal Airways

**DOI:** 10.1101/2025.03.24.644887

**Authors:** Jiaqi Sun, Shisheng Jiang, Hui Sun, Xiaoxiao Xie, Hao Meng, Dong Wang, Hongjie Yao, Jianwei Dai, Hui Zheng, Weijie Guan, Jincun Zhao, Wei Kevin Zhang, Tao Xu, Huisheng Liu

## Abstract

Pulmonary immunity in the human distal respiratory airways is essential for lung function but remains poorly explored, mainly due to limited physiologically relevant models. Here, we develop distal lung organoids containing respiratory airway secretory (RAS) cells, a recently identified epithelial population unique to the distal airways of humans and large mammals, using human pluripotent stem cells (hPSCs). Lineage tracing identified RAS cells as descendants of SOX9^bright^NKX2-1^bright^ progenitors. Single-cell transcriptomics elucidated that RAS cells exhibited a distinct immune-competent phenotype with enriched genes associated with viral host entry and pattern recognition receptor signaling. Functionally, RAS cells infected by respiratory syncytial virus (RSV) exhibited enhanced antiviral activity marked by upregulation of interferon-stimulated genes (ISGs) and complement component 3 (C3). In contrast, bacterial flagellin or *Pseudomonas aeruginosa* (PAO1) triggered a TLR5-driven immune response in RAS cells, inducing complement system activation, a response suppressed by selective TLR5 inhibition. Notably, in chronic obstructive pulmonary disease (COPD), RAS cells displayed dysregulated immune activation, characterized by elevated expression of adaptive immune mediators. Together, our findings identify RAS cells as previously unrecognized sensors and effectors of mucosal immunity in the human distal airways and reveal their roles in viral infection, bacterial stimulation, and chronic inflammatory lung disease, providing potential therapeutic targets for respiratory diseases.

## Introduction

The human respiratory tract orchestrates air exchange while defending against pathogens through specialized epithelial cells ^1,2^. Unlike mice, humans possess distinct distal airway structures, termed respiratory airways (RAs) or terminal and respiratory bronchioles (TRBs), that bridge the conducting airways to the alveolar sacs. These distal structures are essential for efficient respiration ^3,4^ yet are frequent targets of injury in respiratory diseases. Chronic obstructive pulmonary disease (COPD) ^5,6^, pulmonary fibrosis ^7–9^, and viral infections ^10,11^ all provoke inflammation and remodeling of RAs, leading to impaired respiratory function and progressive deterioration. A limited understanding of immune regulation in RAs has hindered the development of effective therapies.

Recent studies have identified unique SCGB3A2-expressing secretory cells (SCs), termed RAS cells or TRB-SCs, in the human distal airways ^12^. These RAS cells serve as progenitors for alveolar type 2 (AT2) cells ^13^. Additional evidence suggests that AT2 cells differentiate into TRB-SCs via an intermediate AT0 state ^14^. Advances in stem cell–based modeling have begun to capture these cells *in vitro*. For example, expandable respiratory airway progenitors (RAPs) have been derived from human pluripotent stem cells and applied to model alveolar differentiation and pulmonary fibrosis^9^, and RAS-like cells have been suggested to arise in certain hPSC lung differentiation cultures^13^. Despite these advances, several fundamental questions remain unresolved. In particular, the developmental origin of RAS cells, their fate plasticity toward airway versus alveolar lineages, and their functional properties—especially with respect to epithelial immune surveillance—have not been systematically examined.

Here we develop a human pluripotent stem cell (hPSC)-derived distal lung organoid system to generate and study RAS cells. These distal lung organoids recapitulate key developmental and structural features of the distal lung epithelium and serve as robust platforms for functional analysis. By combining single-cell transcriptomics, pathogen challenge assays, and analysis of clinical COPD specimens, we show that RAS cells arise from NKX2-1^bright^SOX9^bright^ distal lung progenitors and acquire context-dependent immune competence. Specifically, RAS cells mount robust immune defense responses to respiratory pathogens. In COPD, RAS cells undergo substantial immune-associated transcriptional remodeling characterized by enhanced adaptive immune and cytokine-related pathways. Notably, complement C3 emerges as a convergent immune effector across multiple experimental and disease contexts. Together, these findings identify previously underappreciated immune-responsive functions of RAS cells in the human distal airway and provide new insights into epithelial immune regulation in respiratory infection and chronic lung disease.

## Results

### Robust Generation of Distal Lung Organoids

Human lung development relies on spatially distinct progenitor populations: SOX2^bright^NKX2-1^dim^ proximal lung progenitor cells (p-LPC) drive airway differentiation, while SOX9^bright^NKX2-1^bright^ distal lung progenitor cells (d-LPC) are critical for branching morphogenesis and alveologenesis (Fig. 1A) ^15–19^. To generate SOX9^bright^NKX2-1^bright^ distal lung progenitors from hPSCs, we utilized and modified previously reported protocols for definitive endoderm (DE) and 3D anterior foregut endoderm (AFE) spheroid induction (fig. S1A) ^20^. Inspired by successful colon organoid systems that dissociate hindgut endoderm into Matrigel ^21^, we dissociated AFE spheroids into single cells and embedded them in 3D Matrigel droplets (Fig. 1B). This dissociation step dramatically increased organoid yield (>4-fold) and uniformity (fig. S1, B to E).

**Fig. 1.**
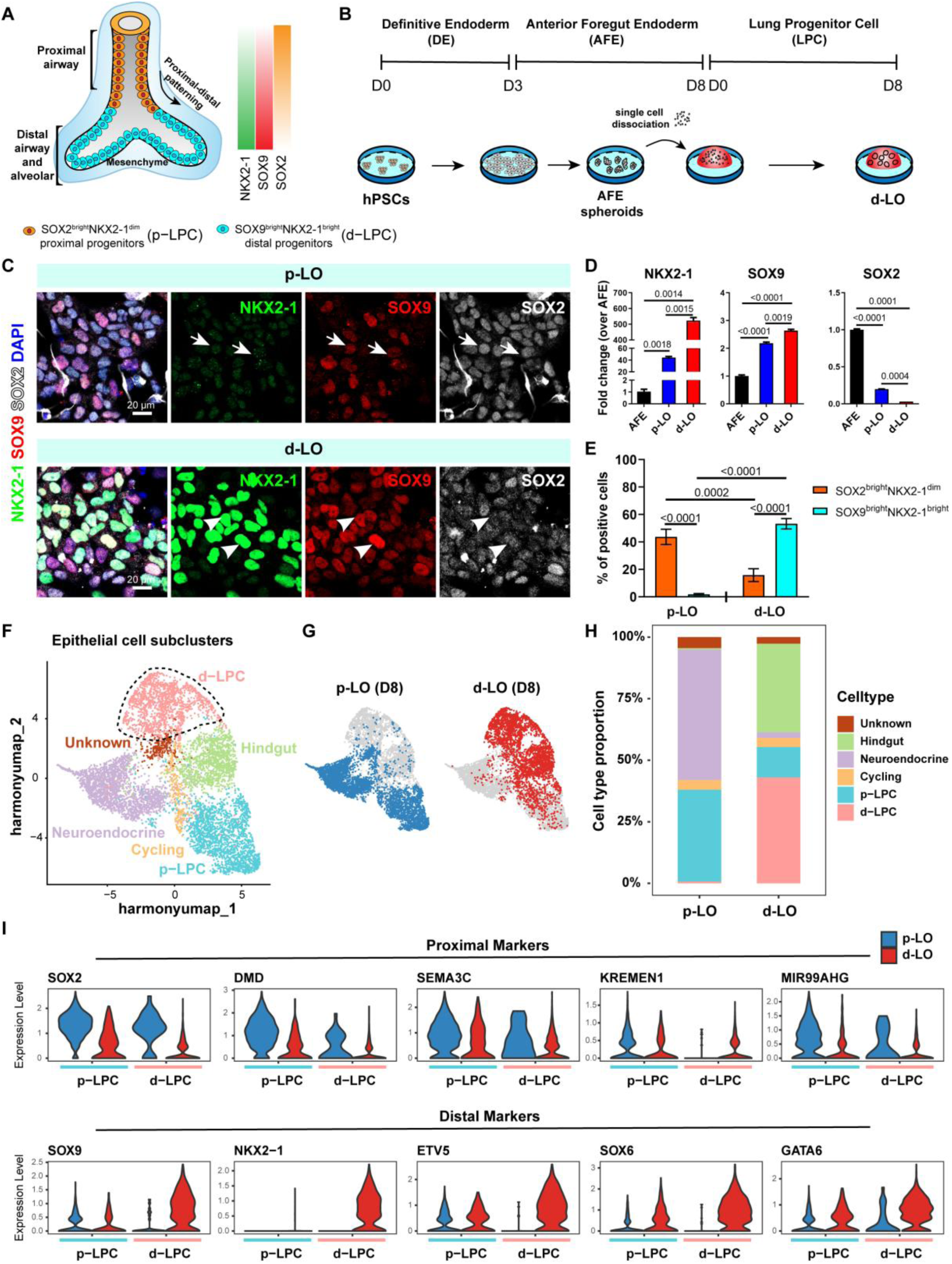
Robust induction of distal lung progenitor cells through AFE spheroid dissociation. (**A**) Schematic of proximal-distal patterning in developing human lung with representative lineage markers. (**B**) Differentiation protocol workflow from human pluripotent stem cells (hPSCs) to distal lung organoids (d-LOs) via dissociation of anterior foregut endoderm (AFE) spheroids. (**C**) Immunofluorescence staining of NKX2-1, SOX2 and SOX9 in proximal (p-LO) and distal (d-LO) lung organoids at day 8 LPC. White arrows denote SOX2^bright^NKX2-1^dim^ cells; arrowheads indicate SOX9^bright^NKX2-1^bright^ cells. Scale bars, 20 μm. Representative images from three biologically independent experiments. (**D**) Quantitative PCR analysis of *NKX2-1*, *SOX9* and *SOX2* expression in AFE spheroids, d-LOs, and p-LOs. Data are presented as mean ± SEM (n=3 biological replicates). *P*-values were calculated using an unpaired two-tailed Student’s *t*-test with Welch’s correction. (**E**) Quantification of the percentage of SOX2^bright^NKX2-1^dim^ (proximal) and SOX9^bright^NKX2-1^bright^ (distal) cells in p-LOs and d-LOs based on immunofluorescence staining analysis. Data are presented as mean ± SEM (n=3 biological replicates). *P*-values were calculated by one-way ANOVA with Tukey’s multiple comparison test. (**F**) UMAP projection annotated with different epithelial cell subcluster identities. (**G**) Individual conditions of epithelial subclusters shown in (**F**). (**H**) Proportional distribution of cell types in d-LOs versus p-LOs. (**I**) Violin plots showing expression levels of proximal and distal markers in p-LPC and d-LPC cell populations derived from d-LOs and p-LOs.

Transcript analysis of lung progenitor markers (*SOX2*, *SOX9*, *NKX2-1*) revealed that dissociation significantly upregulated *NKX2-1* expression (10- to 100-fold), with minimal changes in *SOX9* and *SOX2* levels (Fig. 1D, and fig. S1I). Triple immunostaining confirmed the enrichment of SOX9^bright^NKX2-1^bright^ cells in dissociation-treated cultures, whereas SOX2^bright^NKX2-1^dim^ populations dominated untreated conditions (Fig. 1, C and E). Flow cytometry further validated elevated NKX2-1 protein levels in dissociated organoids (fig. S1, F to H). Aligning with embryonic lung gene expression patterns ^15^, we designated organoids from untreated conditions as proximal lung organoids (p-LOs) and those from dissociated cultures as distal lung organoids (d-LOs). The protocol consistently generated d-LOs across multiple hPSC lines, underscoring its robustness and broad applicability (fig. S1I).

### Characterization of d-LOs

To validate the distal identity of d-LOs, we performed single-cell RNA sequencing (scRNA-seq) on 10,815 cells from p-LOs and d-LOs at day 8. Our analysis identified three cellular compartments: epithelial, mesenchymal, and neural cells (fig. S1, J and K). Compared with p-LOs, d-LOs exhibited a reduced proportion of neural cells and an increased mesenchymal compartment (fig. S1L). Prior studies have shown that mesenchymal cells can promote bud tip maintenance and multipotency^22^, suggesting a potential supportive role for the expanded mesenchymal population observed in d-LOs.

We then extracted and reclustered the epithelial cells for a more detailed investigation (Fig. 1, F and G, and fig. S1M). Six cell clusters were annotated based on their distinct molecular features: d-LPC, p-LPC, cycling, neuroendocrine, hindgut, and an unknown population. The d-LPC and p-LPC clusters comprised lung-fated progenitors distinguished by differential expression of *NKX2-1* together with *SOX2* and *SOX9*. Cycling cells were marked by high expression of proliferation-associated genes including *TOP2A*, *CDK1*, and *UBE2C*. In addition, we identified a cluster of STMN2⁺SST⁺ neuroendocrine-like cells, as well as a discrete population expressing SOX17 and CDX2, which we annotated as hindgut. Notably, the d-LPC cluster was abundantly generated in d-LOs (43.01% in d-LOs and 0.71% in p-LOs; Fig. 1H), while p-LPC was enriched in p-LOs (12.33% in d-LOs and 37.29% in p-LOs; Fig. 1H), corroborating gene and protein expression data.

Benchmarking against human fetal lung datasets ^15^ further confirmed this proximal-distal divergence (Fig. 1I and fig. S1P): p-LPC in p-LOs expressed proximal markers (e.g., *SOX2*, *DMD*, *SEMA3C*, *KREMEN1*, and *MIR99AHG*), while the d-LPC cluster in d-LOs upregulated distal regulators (e.g., *SOX9*, *ETV1*, *SOX6*, and *GATA*). Notably, ETV1 and SOX6 are human development-specific transcription factors involved in distal lung patterning ^15^. Consistent results were obtained when benchmarking p-LPCs and d-LPCs against additional human embryonic lung atlases spanning broader developmental windows^23,24^, further supporting their assignment as proximal and distal progenitor populations (fig. S1, Q-S).

Gene Ontology (GO) analysis of differentially expressed genes (DEGs) within the p-LPC and d-LPC clusters reinforced their proximal and distal identities (fig. S1, N and O). Specifically, p-LPC signature genes were involved in cilium organization and assembly, while d-LPCs were enriched for genes associated with epithelial tube morphogenesis and branching morphogenesis, with high expression of *NKX2-1*, *SOX9*, *WNT7B* and low expression of *SFTPC*, mirroring 7–8 post-conception weeks (pcw) fetal lung bud tips ^19^. Collectively, these data strongly suggest that the d-LPC population in d-LO closely recapitulates the transcriptional profile and functional characteristics of early human lung bud tip cells.

### Differentiation potential of d-LOs

To assess the fate potential of d-LOs in our system, we cultured p-LOs and d-LOs either in airway (DCIK) ^25^ or in alveolar (DCIK+CSB) media ^26^ (fig. S2A). We found that p-LO-derived airway organoids (p-AWO) expressed higher levels of proximal airway genes like *TP63* and *MUC5AC*, while d-LO-derived airway organoids (d-AWO) showed elevated expression of distal airway lineage genes, including *SCGB3A2* (a distal club cell marker), and alveolar lineage markers (*SFTPB*, *AGER*, *PDPN*, and *LAMP3*) (fig. S2B). For alveolar induction, only d-LOs cultured in DCIK+CSB media (d-ALO) exhibited robust alveolar differentiation, marked by high levels of *SFTPB*, *SFTPC*, *NKX2-1*, and *LAMP3*, consistent with the requirement of NKX2-1 for distal lung development. Notably, *SCGB3A2* was modestly expressed in d-ALO. Immunostaining further confirmed that d-LOs readily differentiated into distal airway and alveolar organoids, with abundant SCGB3A2 and SFTPB co-expressing cells (Fig. 2A), which were rarely detected in p-AWO (Fig. 2B and fig. S2C). Collectively, these findings suggest that p-LOs predominantly generate proximal airway organoids, while d-LOs uniquely model human distal airway and alveoli (fig. S2D).

**Fig. 2.**
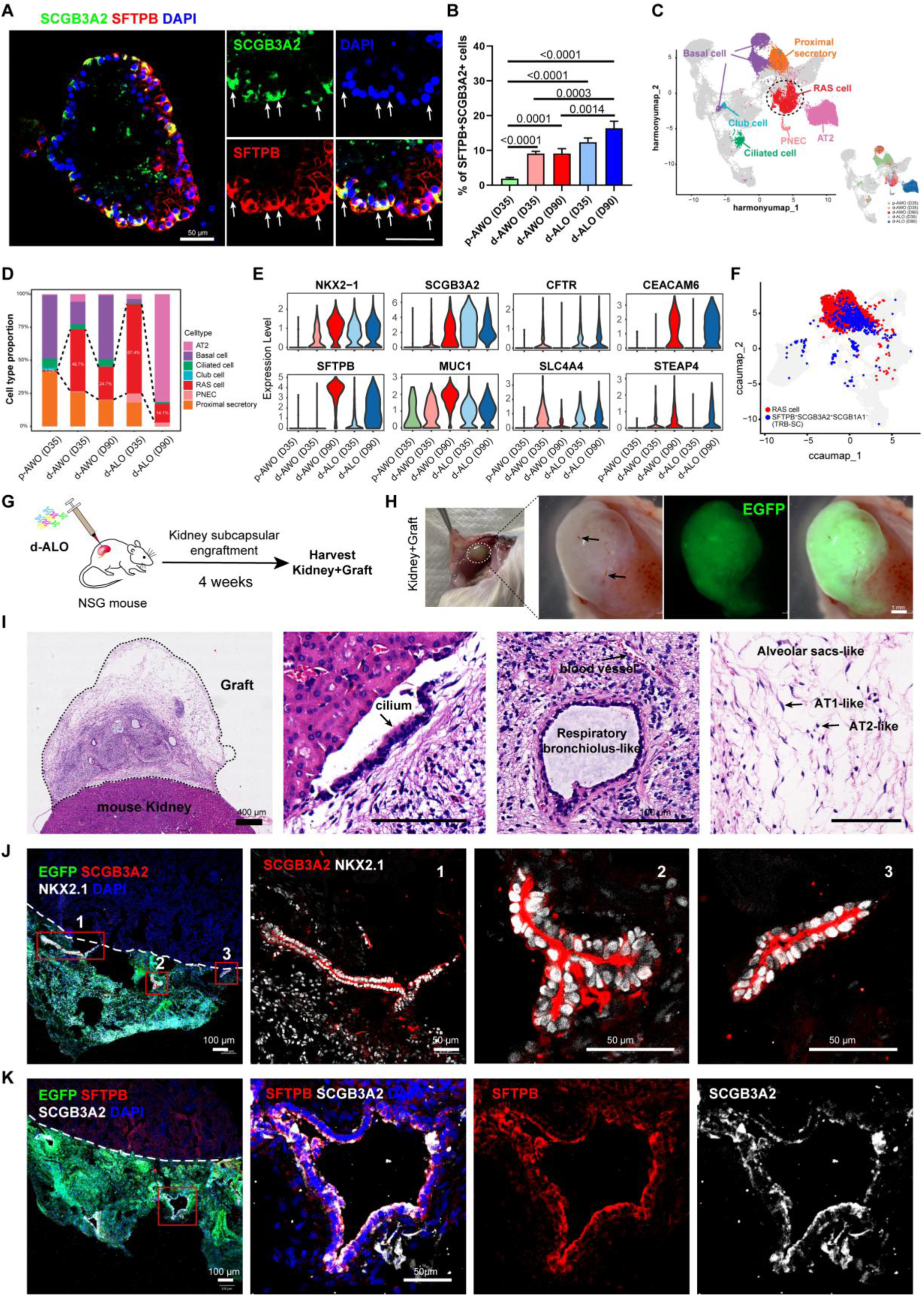
scRNA sequencing and transplantation models delineate the marker expression and in vivo maturation of RAS cells. (**A**) Immunofluorescence staining of SCGB3A2 (green) and SFTPB (red) in d-ALOs. Arrows indicate SCGB3A2^+^SFTPB^+^ co-expressing RAS cells. Scale bars, 50 μm. (**B**) Quantification of the percentage of SFTPB^+^SCGB3A2^+^ cells in p-LO- and d-LO-derived organoids at different time points (D35, D90). Results are representative of 3 independent experiments. Data are presented as mean ± SEM. *P* values are derived from one-way ANOVA with Tukey’s multiple comparison test. (**C**) UMAP embeddings colored by annotated cell types from single-cell RNA sequencing analysis of p-LO- and d-LO-derived organoids (inset shows embeddings colored by differentiation time/media conditions). (**D**) Proportional distribution of lung lineage cell types in d-LO-and p-LO-derived organoids. (**E**) Violin plots illustrating the expression of *NKX2-1*, *SCGB3A2*, *CFTR*, *CEACAM6*, *SFTPB*, *MUC1*, *SLC4A4*, and *STEAP4* within the RAS cell cluster across different time/media conditions. (**F**) UMAP projection showing the overlap between the annotated RAS cell cluster (red) and the SFTPB^+^SCGB3A2^+^SCGB1A1^-^ TRB-SC signature (blue). (**G**) Workflow for subrenal capsule transplantation of d-ALOs into NSG mice. (**H**) Macroscopic images of transplanted d-ALOs (green) at 4 weeks post-engraftment. Arrows indicate vasculature. Scale bar, 1 mm. (**I**) Hematoxylin and eosin (H&E) staining of d-ALO grafts at 4 weeks post-transplantation. Scale bar, 400 µm (left), 100 μm (middle/right). (**J**) Immunofluorescence imaging of transplanted d-ALOs at 4 weeks post-engraftment. Triple staining for SCGB3A2 (red), EGFP (green), and NKX2-1 (grey) (left panel). Scale bars, 100 μm (wide-field images), 50 μm (magnified images). Representative images from three biologically independent experiments. (**K**) Immunofluorescence imaging of transplanted d-ALOs at 4 weeks post-engraftment. Triple staining for SFTPB (red), EGFP (green), and SCGB3A2 (grey). Scale bars, 100 μm (wide-field images), 50 μm (magnified images). Representative images from three biologically independent experiments.

Additionally, d-LO-derived organoids demonstrated functional properties, including CFTR-dependent forskolin-induced swelling (fig. S3, A and B) and BODIPY-labeled phosphatidylcholine uptake and secretion (fig. S3, E and F). Beating cilia were visible (movie S1) in apical-out d-AWO released from Matrigel ^27^ (fig. S3C). Transmission electron microscopy (TEM) analysis of d-ALOs revealed lamellar body-like organelles (fig. S3D). In summary, these results indicate that d-LOs can efficiently generate distal airway and alveolar organoids with functional and structural features characteristic of mature lung tissue.

### RAS cells in d-LO-derived organoids

To further investigate the unique cell types in d-LO-derived organoids, we performed scRNA-seq on p-AWO (D35), d-AWO (D35 and D90), and d-ALO (D35 and D90) (figs. S2, A, E and F). Cell populations were annotated using differentially expressed genes and correlated with established cell signatures from prior studies ^28^ (fig. S2, G and H, Supplementary Table 4). Both p-AWO and d-AWO showed abundant basal and proximal secretory cells, alongside smaller populations of club cells, ciliated cells, and pulmonary neuroendocrine cells (PNECs). AT2 cells were prominently enriched in d-ALO (D90) (Fig. 2, C and D). Notably, within d-LO-derived organoids, we identified a distinct population of SCGB3A2^+^SFTPB^+^SFTPC^-^ cells, which we termed respiratory airway secretory cells (RAS cells), consistent with a distal respiratory airway identity.

To assess the extent to which our organoids recapitulated the cellular landscape of the human distal airways, we compared our dataset with a published single-cell atlas of human distal airways from Murthy et al.^14^. Reference-based mapping using Seurat MapQuery demonstrated that our organoids captured the major epithelial populations described in human distal airways, including AT1, AT0, TRB-SCs, Distal-BC-1, Distal-BC-2, basal cells, neuroendocrine cells, and ciliated cells (fig. S3, G and H). We further performed an integrated analysis, which revealed extensive co-localization of organoid-derived cells with their in vivo counterparts in shared transcriptional space, supporting a high degree of transcriptional similarity at the lineage level (fig. S3, I to K).

The unique RAS cluster was positioned between the d-LPC, proximal secretory, PNEC, and AT2 clusters on the UMAP embedding (Fig. 2C and fig. S2G), exhibited a specific gene expression profile, including markers such as *MUC1*, *CFTR*, *CEACAM6*, *SLC4A4*, and *STEAP4* (Fig. 2E), with MUC1 and CEACAM6 serving as cell surface markers for these specialized secretory cells ^13,29^. To quantify the abundance of RAS cells across organoid conditions and developmental stages, we analyzed their proportions in our scRNA-seq datasets relative to both all cells and the epithelial compartment (Fig. 2D and fig. S2I). In p-AWOs at day 35, RAS cells were rare, comprising approximately 0.6% of all cells and ∼1.2% of epithelial cells. In contrast, RAS cells were markedly enriched in distal-patterned organoids. In d-AWOs (D35), they accounted for ∼8.2% of all cells and 46.7% of epithelial cells, whereas in d-ALOs (D35) they represented ∼17.8% of all cells and 67.4% of epithelial cells. At later stages (D90), RAS cells remained detectable but at reduced relative abundance, comprising ∼9.1% of all cells and 24.7% of epithelial cells in d-AWOs, and ∼10.2% of all cells and ∼14.1% of epithelial cells in d-ALOs (Fig. 2D and fig. S2I). Notably, a substantial fraction of cells annotated as RAS in our dataset preferentially mapped to the TRB-SC compartment (fig. S3H). Moreover, integration analysis demonstrated that organoid-derived RAS cells extensively overlapped with primary human TRB-SCs, supporting their close transcriptional resemblance (Fig. 2F).

To further assess distal airway potential *in vivo*, d-ALOs were transplanted under the kidney capsule of immunodeficient NSG mice. After 4 weeks, the grafts developed organized tubular and saccular structures (Fig. 2, G to I). The tubular structures were lined by NKX2-1^+^SCGB3A2^+^ epithelium, which also expressed SFTPB (Fig. 2, J and K), mirroring the distal characteristics of human fetal lung tissue at 19–20-pcw ^14^. Collectively, these findings underscore the unique differentiation potential and functional relevance of d-LO-derived organoids in modeling human respiratory airways.

### A d-LPC–RAS cell relationship

Research shows that SCGB3A2-expressing cells emerge as early as 5 pcw in fetal lungs ^24^. To explore the developmental trajectory of RAS cells, we integrated scRNA-seq data from d-LOs and d-LO-derived organoids (fig. S4, A to C). Trajectory inference analysis using Monocle3 revealed a strong cell trajectory originating from the d-LPC cluster into RAS cell cluster (Fig. 3, A to C). Additional trajectory analyses using Slingshot and diffusion pseudotime yielded highly similar results, independently supporting the developmental progression from d-LPCs toward RAS cells (fig. S5). The gene expression changes along the d-LPC–RAS cell trajectory highlighted several genes known to be involved in secretory and alveolar differentiation, including *SCGB3A2*, *SFTPB* and *HOPX* (Fig. 3, D and E). Additionally, transcription factors like *AHR*, *GRHL1*, and *SOX2*, identified through pseudotime analysis, showed strong correlations with primary distal airway secretory cells^14^.

**Fig. 3.**
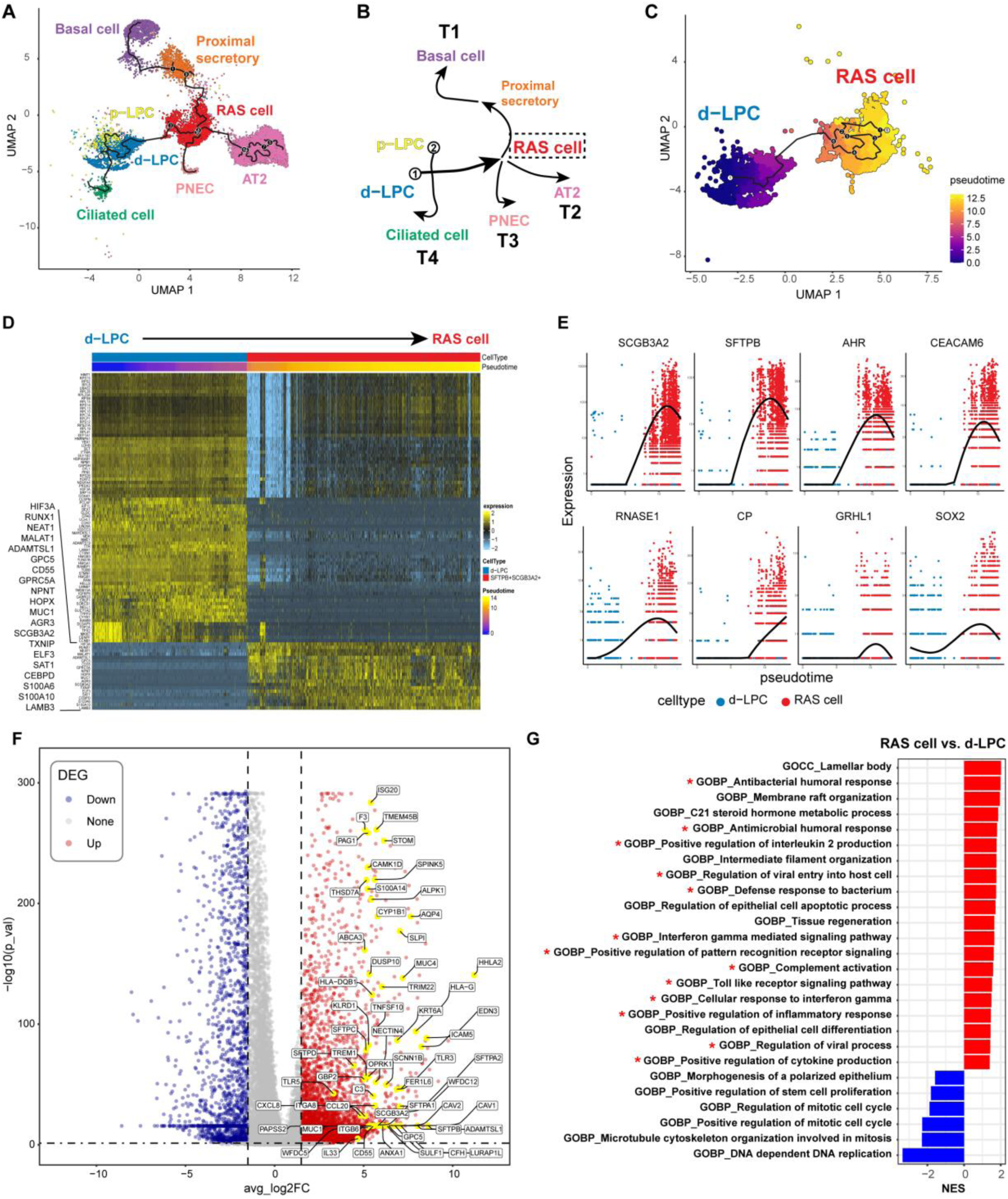
Developmental trajectories and functional characterization of RAS cells. (**A**) Pseudotemporal trajectory reconstruction of epithelial subtypes using Monocle3. (**B**) Schematic summarizing developmental lineages inferred from pseudotemporal analysis in (**A**). (**C**) Differentiation trajectory connecting d-LPCs to RAS cells. (**D**) Heatmap of gene expression dynamics along the pseudotemporal trajectory from d-LPCs to RAS cells (pseudotime scale above). (**E**) Smoothed expression patterns of select lineage-associated genes during d-LPC-to-RAS cell differentiation. (**F**) Volcano plot of differentially expressed genes (DEGs) between d-LPCs and RAS cells. (**G**) Gene set enrichment analysis (GSEA) of DEGs from (**F**), ranked by normalized enrichment score (NES).

Further analysis of the integrated dataset demonstrated pseudotemporal relationships between RAS cells and basal, PNEC, and AT2 cells (Fig. 3, A and B, and fig. S4, D to G), indicating that RAS cells may function as progenitors for human distal lung development and regeneration. Trajectory inference also indicated a developmental link between p-LPC and ciliated cells (fig. S4H). Together, these in silico predictions suggest that d-LPC acts as a putative progenitor for the RAS cell lineage, which can subsequently give rise to basal, PNEC, and AT2 cells.

### RAS cells exhibit immune functions

In addition to the developmental progression inferred from trajectory analysis, differential gene expression analysis between d-LPCs and RAS cells revealed increased expression of multiple immune-associated genes in RAS cells, including *RUNX1*, *CD55*, *SCGB3A2*, *TXNIP*, *CEBPD*, *S100A6*, and *S100A10*, as well as mucins (*MUC1*, *MUC4*), surfactant proteins (*SFTPB*, *SFTPD*), cytokines (*CXCL8*, *CCL20*), and cell adhesion molecules (*ICAM5*, *ITGB6*) (Fig. 3F). Gene set enrichment analysis (GSEA) of the DEGs revealed significant enrichment of immune-associated pathways, such as antibacterial responses, regulation of viral entry into host cells, and positive regulation of pattern recognition receptor (PRR) signaling (Fig. 3G). These findings align with recent single-cell profiling data from the human distal lung, indicating through pathway enrichment analysis that TRB-SCs may possess immunomodulatory functions^14^.

To further compare these programs across epithelial populations, we calculated module scores for the major immune pathways identified from the RAS-versus-dLPC analysis. RAS cells showed the highest enrichment of pattern-recognition receptor signaling, cytokine-production, complement-activation, and interferon-response programs among distal epithelial populations (Fig. S6), indicating that immune-sensing and inflammatory-response programs are particularly prominent in RAS cells.

Together, these results establish a broad spectrum of immune-related transcriptional activity in RAS cells, emphasizing their pivotal role in respiratory immune defense.

### RAS cells mount broad interferon-driven and inflammatory responses to RSV infection

Given the clinical relevance of RSV infection in distal airways in vulnerable individuals—particularly as a leading cause of bronchiolitis and pneumonia ^30^, we therefore evaluated RSV susceptibility using our distal organoid model (Fig. 4A).

**Fig. 4.**
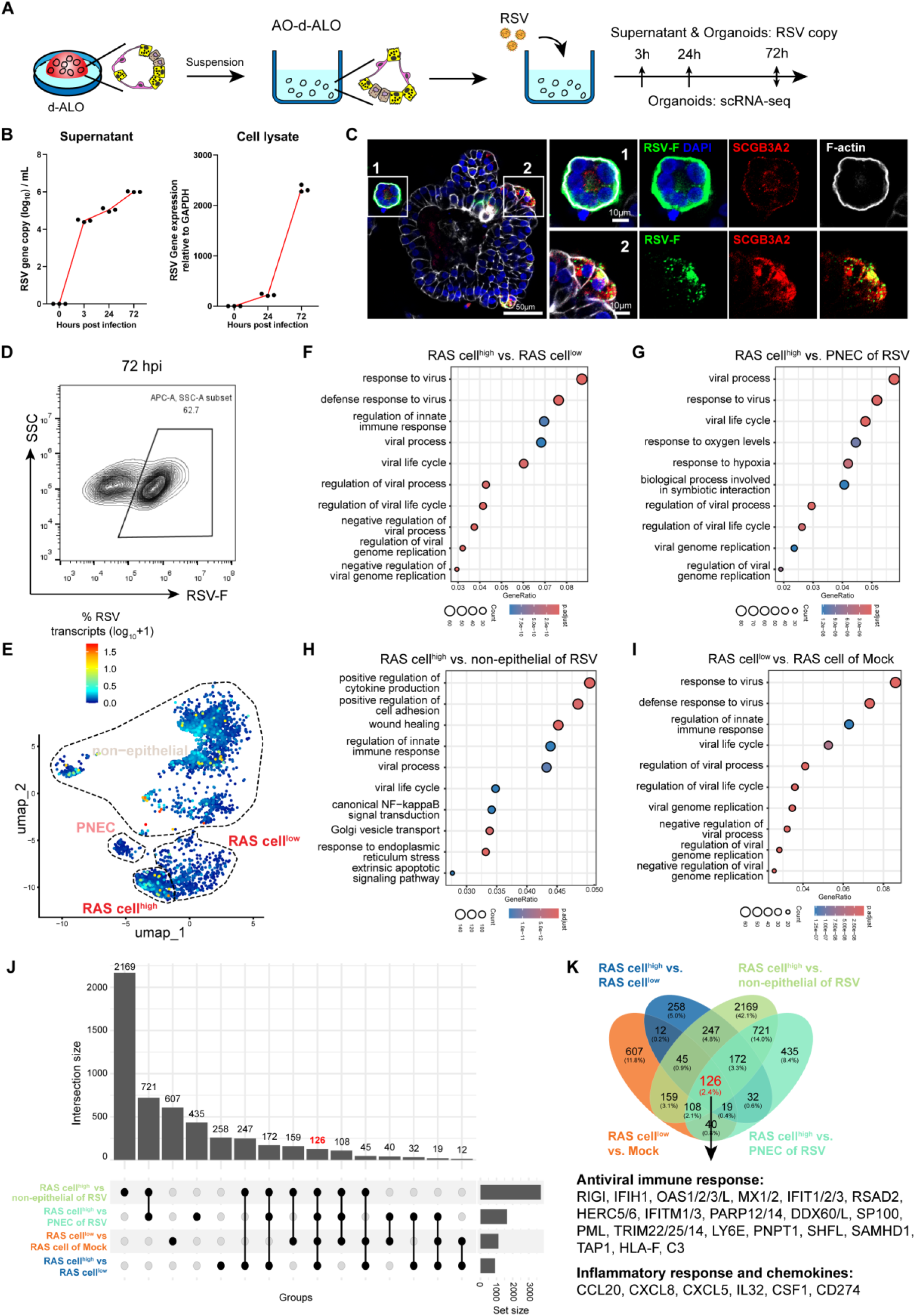
Enhanced antiviral activity of RAS cells during RSV infection. (**A**) Schematic illustration of the procedure for RSV infection in D40 d-ALOs. (**B**) RSV replication kinetics in d-ALOs quantified by RT-qPCR of viral RNA in supernatants (left) and cell lysates (right) at indicated time points. Each dot represents one well (n = 3 biological replicates). (**C**) Morphological changes in d-ALOs post-RSV infection. Left: overview; right: magnified regions showing cytopathic effects. Scale bars, 50 μm (overview), 10 μm (magnified images). Representative images from three independent experiments. (**D**) Infection efficiency in d-ALOs assessed by flow cytometry analysis of RSV-F glycoprotein expression at 72 hpi (MOI= 0.1). (**E**) UMAP projection of single-cell transcriptomes colored by the percentage of viral transcripts per cell. (**F** to **I**) Gene Ontology (GO) enrichment analysis of differentially expressed genes (DEGs) identified in the following comparisons: (**F** to **G**) RAS cell^high^ versus other clusters in infected organoids and (**I**) RAS cell^low^ versus Mock. (**J** to **K**) Venn diagrams illustrating shared DEGs between RAS cell^high^ versus other clusters (Infected), and RAS cell^low^ versus the same cluster in mock.

RSV exhibited robust replication kinetics at a low multiplicity of infection (MOI = 0.1), as quantified in both supernatant and cell lysates (Fig. 4B). By 72 hpi, 62.7% of organoid cells were positive for RSV-F protein (Fig. 4D). Morphological analysis of infected d-ALOs revealed significant epithelial malformations, including cytoskeletal reorganization and syncytia formation (Fig. 4C), mirroring clinical hallmarks of RSV infection ^31^. Notably, SCGB3A2-positive cells were extensively targeted by RSV, as confirmed by co-staining of SCGB3A2 and RSV-F. Infected organoids exhibited substantial upregulation of inflammatory mediators (*CXCL8*, *CXCL10*, *TNF-a*, *IL-6*) (fig. S7A), contributing to immune cell recruitment and airway inflammation.

To further explore the cellular response, we performed scRNA-seq following RSV challenge (fig. S7B). The scRNA-seq data revealed widespread infection, with viral transcripts detected in nearly all cells (Fig. 4E). However, viral gene expression was unevenly distributed, with most cells showing low viral transcript levels and two distinct clusters exhibiting significantly higher expression—one within RAS cells (RAS cell^high^) and another among non-epithelial cells.

We next conducted DEGs analysis between RAS cell^high^ and other infected clusters, as well as between RAS cell^low^ and mock-treated counterparts. GO analysis revealed consistent enrichment of terms such as “response to virus” and “viral life cycle” across clusters (Fig. 4, F to I). A set of 126 overlapping genes was identified across comparisons (Fig. 4, J and K), with significant enrichment of antiviral innate immune response, NF-κB signaling, and antigen-presentation pathways (fig. S7, C and D). Notably, these RSV-responsive genes were most strongly induced in RAS^high^ cells (fig. S8, A and B), suggesting that the RAS cells actively participate in pathogen sensing and that cells with higher viral burden mount a more robust antiviral response.

We observed elevated expression of PRRs in RAS cell^high^, including *RIG-I* (DDX58) and *MDA5* (IFIH1), indicating robust viral RNA sensing and interferon pathway activation. Consistently, numerous interferon-stimulated genes (ISGs), chemokines, and inflammatory mediators were markedly induced (Fig. 4K). Genes involved in antigen processing and immune regulation, including *TAP1*, *CD274*, and *HLA-F*, were also upregulated. Notably, complement component C3 was consistently elevated.

To further compare epithelial responses to RSV infection, infected cells were projected onto an integrated distal lung epithelial reference atlas, identifying RAS, AT2, basal, club, ciliated, proximal secretory, and PNEC populations. The proportion of RAS cells was largely preserved following infection compared (fig. S8, C-D). Notably, the RAS Cell^high^ subset, together with ciliated cells, exhibited higher levels of RSV transcripts (Fig. S8, E-G). We next calculated a signature score based on the 126 RSV-responsive genes identified above. Although antiviral responses were observed across multiple epithelial populations, RAS Cell^high^ displayed the strongest enrichment of the RSV-response signature and significantly higher scores than AT2, ciliated cells, and most other epithelial populations (Fig. S8, H-J). Together, these findings indicate that RAS cells represent a major epithelial responder to RSV infection within the distal airway epithelium.

To determine whether this immune activation was accompanied by proliferative changes, we performed scRNA-seq–based cell-cycle analysis. RAS cells did not undergo proliferative expansion following infection, but instead exhibited relative cell-cycle arrest, with a marked reduction in the proportion of cells in S and G2/M phases compared to mock control (S+G2/M ∼33% in RSV conditions versus ∼46% in mock) (fig. S7, E and F).

Consistently, expression of canonical proliferation-associated genes, including *PCNA*, *MCM2*–*6*, and *CCNB1*, was reduced in RSV-infected RAS cells (fig. S7G). These results indicate that the infection-induced immune gene upregulation observed in RAS cells is not associated with increased proliferation, but rather reflects activation of host-defense programs in a relatively non-proliferative state.

To provide an independent human-lung benchmark for these antiviral programs, we analyzed a publicly available single-cell transcriptomic dataset from COVID-19 patients^32^. Re-annotation of epithelial populations identified an airway club cell population sharing key transcriptional features with RAS cells, including expression of *SCGB3A2*, *CFTR* and *SFTPB* (Fig. S7H). Of the 126 RSV-responsive genes identified in our organoid dataset, 121 were detected in the COVID-19 dataset, and a substantial subset was similarly upregulated in airway club cells from COVID-19 lungs compared with controls (Fig. S7I).

Together, these findings support the physiological relevance of the immune activation states observed in our RSV infection model and suggest that RAS cells possess prominent immune-responsive properties during viral infection, including activation of interferon signaling, inflammatory pathways, and local complement programs.

### RAS cells display antimicrobial responses via activation of TLR5

GSEA revealed significant enrichment of Toll-like receptor (TLR) signaling pathway genes in RAS cells (Fig. 3G), a pathway central to respiratory tract host defense ^33^. Notably, TLR5, a flagellin sensor, was highly expressed in RAS cells (fig. S9, A and B). This expression pattern aligned with scRNA-seq data from early human fetal lung tissue^34^, in which a SCGB3A2^+^SFTPB^+^CFTR^+^ subpopulation expresses TLR5 at higher levels than other epithelial cells (fig. S9, C and D). This suggests that, although organoid-derived RAS cells resembled adult TRB-SCs at the transcriptomic level, their TLR5 expression pattern reflected a developmental distal airway epithelial state. Immunostaining further confirmed robust TLR5 protein expression in RAS cells within organoids (Fig. 5, A and B).

**Fig. 5.**
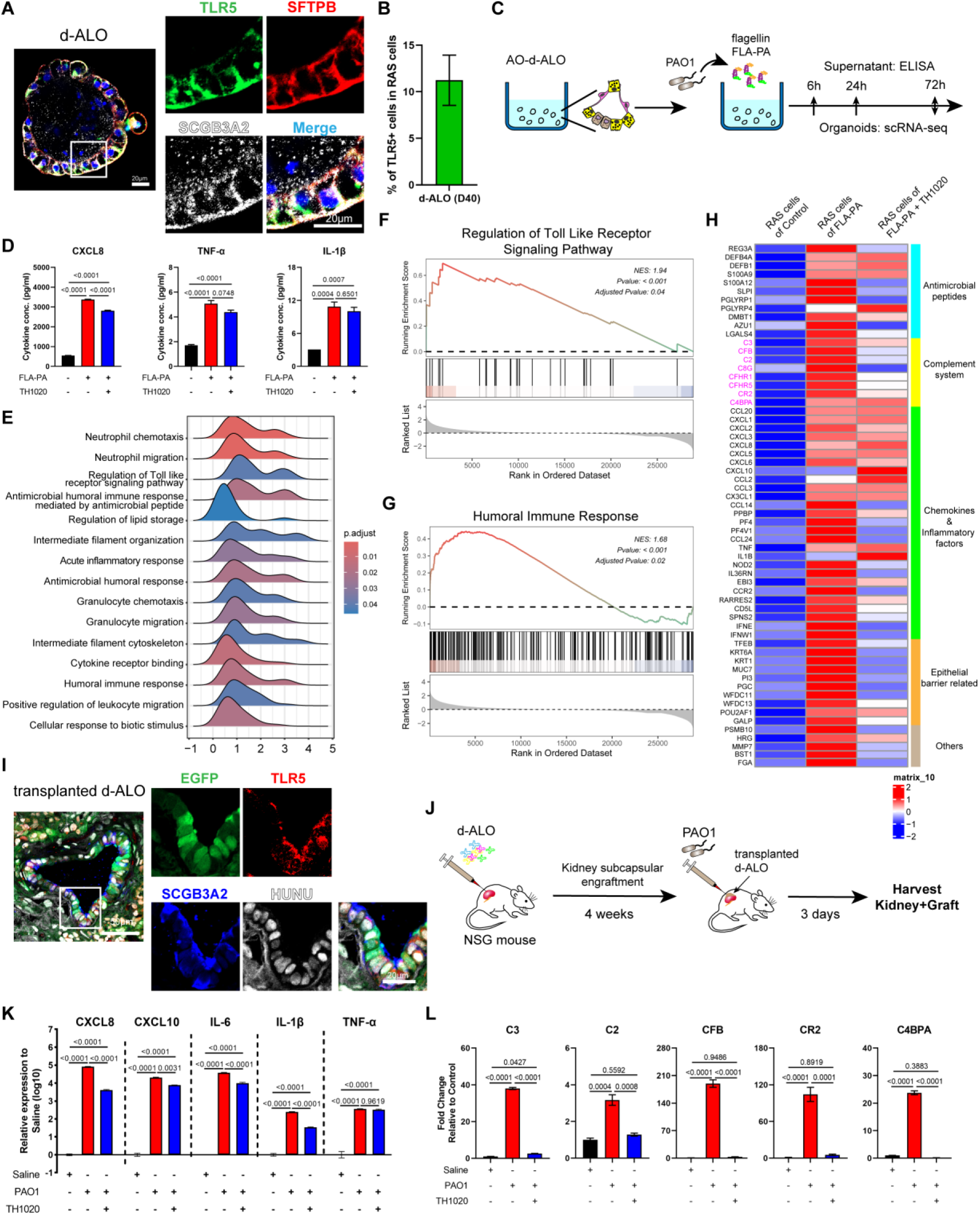
TLR5 activation mediates microbial-induced immune responses in RAS cells. (**A**) Whole-mount immunofluorescence staining for TLR5 (green) and RAS cell markers (SFTPB, red and SCGB3A2, grey) in d-ALO. Nuclei counterstained with Hoechst (blue). Scale bars, 20 μm. Representative of three independent experiments. (**B**) Quantification of the percentage of TLR5^+^ cells within the RAS cell population in d-ALOs (Day 40). Results are representative of 3 independent experiments. Data are presented as mean ± SEM. (**C**) Schematic illustration of the procedure for FLA-PA treatment in Day 40 d-ALOs. (**D**) Secretion of cytokines/chemokines (CXCL8, CXCL10, IL-6, IL-1β, TNF-α) from d-ALOs stimulated with 1 μg/ml FLA-PA protein for 72 hours, with or without 1 μM TLR5 inhibitor TH1020. (**E**) Top 15 GO terms enriched in DEGs between FLA-PA-treated RAS cells and control RAS cells. (**F**, **G**) Gene Set Enrichment Analysis (GSEA) plots showing up-regulation of the TLR signaling pathway (**F**) and the humoral immune response (**G**) in FLA-PA-treated RAS cells compared to control RAS cells. (**H**) Expression profiles of genes enriched in the “humoral immune response” pathway (from GSEA) across control, FLA-PA-treated, and FLA-PA+TH1020-treated RAS cells. (**I**) Immunofluorescence co-staining for TLR5 (red), SCGB3A2 (blue), and human nuclear antigen (HUNU, white) in transplanted d-ALOs at 4 weeks post-engraftment in NSG mice. Scale bars, 50 μm (low-magnification images), 20 μm (magnified images). Representative of three independent transplants. (**J**) Schematic illustration of the procedure for PAO1 infection in transplanted d-ALOs. (**K**) Quantitative PCR analysis of inflammatory mediators in transplanted d-ALOs stimulated with PAO1 lysate for 72 hours, with or without 1 μM TH1020 treatment. (**L**) Quantitative PCR analysis of complement-related genes in PAO1 lysate-stimulated transplanted d-ALOs with or without 1 μM TH1020 (72 h). (**D**,**K**,**L**) Data are presented as mean values ± SEM. *P*-values were calculated by one-way ANOVA with Tukey’s multiple comparison test (n=3 biological replicates).

To assess the functional role of TLR5 activation in RAS cells in response to flagellin, we treated our distal organoids with *Pseudomonas aeruginosa* (PAO1)-derived flagellin (FLA-PA) (Fig. 5C). Stimulation with FLA-PA significantly promoted the secretion of inflammatory mediators, including IL-6, CXCL8, CXCL10, and TNF-α (fig. S9E). This response was modestly attenuated by the TLR5-specific inhibitor TH1020 (Fig. 5D).

To further investigate the transcriptional response of RAS cells to TLR5 activation, we performed scRNA-seq on organoids treated with FLA-PA, with or without TH1020 (fig. S9, F to H). GSEA revealed significant induction of 29 canonical pathways in FLA-PA-treated RAS cells compared to controls (fig. S10, A to F and Supplementary table S2). Among the top 15 enriched pathways, the majority were immune-related, including neutrophil chemotaxis/migration, antimicrobial peptide-mediated responses, Toll-like receptor signaling, and the humoral immune response (Fig. 5, E to G and fig. S9I), indicating robust activation of innate immune programs upon TLR5 stimulation.

To determine whether this immune activation was accompanied by changes in cell proliferation, we performed a cell-cycle analysis using scRNA-seq data. FLA-PA stimulation did not result in a marked increase in RAS cell proliferation at 72 h (fig. S9, J to L). Strikingly, inhibition of TLR5 signaling by TH1020 led to a modest increase in proliferative activity, as reflected by an elevated proportion of cells in S and G2/M phases and upregulation of cell-cycle–associated genes. These findings indicate that TLR5 activation in RAS cells primarily drives transcriptional activation of innate immune programs rather than proliferative expansion.

Under the GO term *humoral immune response*, 61 genes were significantly upregulated in RAS cells following FLA-PA exposure, encompassing chemokines, antimicrobial effectors, and epithelial-associated transcripts (Fig. 5H). Notably, multiple complement-related genes—including *C2*, *C3*, *CFB*, *CFHR1*, *CFHR5*, *C4BPA*, *C8G*, and *CR2*—were markedly induced. The transcriptional activation of complement components is likely to facilitate opsonization and microbial clearance at the mucosal surface ^35^. Importantly, co-treatment with TH1020 substantially abolished the upregulation of these complement genes (Fig. 5H), while chemokine and antimicrobial gene expression persists to some extent. This suggests that the complement response is strictly TLR5-dependent, whereas the induction of chemokines and antimicrobial genes by FLA-PA may involve additional pathways, such as NOD2 signaling, in parallel with TLR5 activation. Furthermore, we observed a similar expression pattern in transplanted d-ALOs stimulated with flagellated bacteria PAO1 (Fig. 5, I to L).

To examine whether this immune activation was specific to RAS cells, we performed additional cell-type–resolved analyses using our scRNA-seq dataset following FLA-PA treatment. GSEA revealed that Toll-like receptor signaling and humoral immune response pathways were not robustly enriched in basal cells, ciliated cells, PNECs, cycling cells, or mesenchymal populations (fig. S10J). Among non-lung epithelial cells, the humoral immune response pathway showed only modest enrichment (adjusted *P* value= 0.01), whereas TLR signaling was not significantly enriched. These results indicate that the transcriptional response to flagellin stimulation is predominantly observed in RAS cells within the organoid cultures. Consistent with this, purified AT2 cells isolated using the HT2-280 antibody did not exhibit immune activation following FLA-PA treatment (fig. S10, G to I).

These results reveal that RAS cells not only sense microbial threats via TLR5 but also mount a tiered defense response involving chemokine secretion, antimicrobial peptides, and complement activation. The unique TLR5 sensitivity of the complement module highlights a regulatory checkpoint linking epithelial sensing to immune effector coordination in distal airways.

### COPD is associated with atypical immune activity of RAS cells

We next investigated whether similar or distinct immune programs are engaged by RAS cells in chronic disease settings, where infection and inflammation persist. To this end, we analyzed single-cell transcriptomic data from the terminal airway regions of COPD patients ^13^ (fig. S11, A to C). RAS cells were detected and clustered in all the analyzed COPD patient and healthy control samples from the public dataset. RAS cells from COPD lungs exhibited elevated expression of *TLR2*, *TLR3* and *TLR5* (Fig. 6A), accompanied by enhanced TLR signaling activity (fig. S11D), suggesting heightened epithelial pathogen sensing. Clinical evidence highlights the pathological relevance of PAO1 lung colonization in COPD ^36^. Immunostaining revealed increased co-expression of SFTPB, SCGB3A2, and TLR5 in infected COPD tissues compared to controls, along with robust expression of chemokines, implicating RAS cells in driving localized inflammation (fig. S11E).

**Fig. 6.**
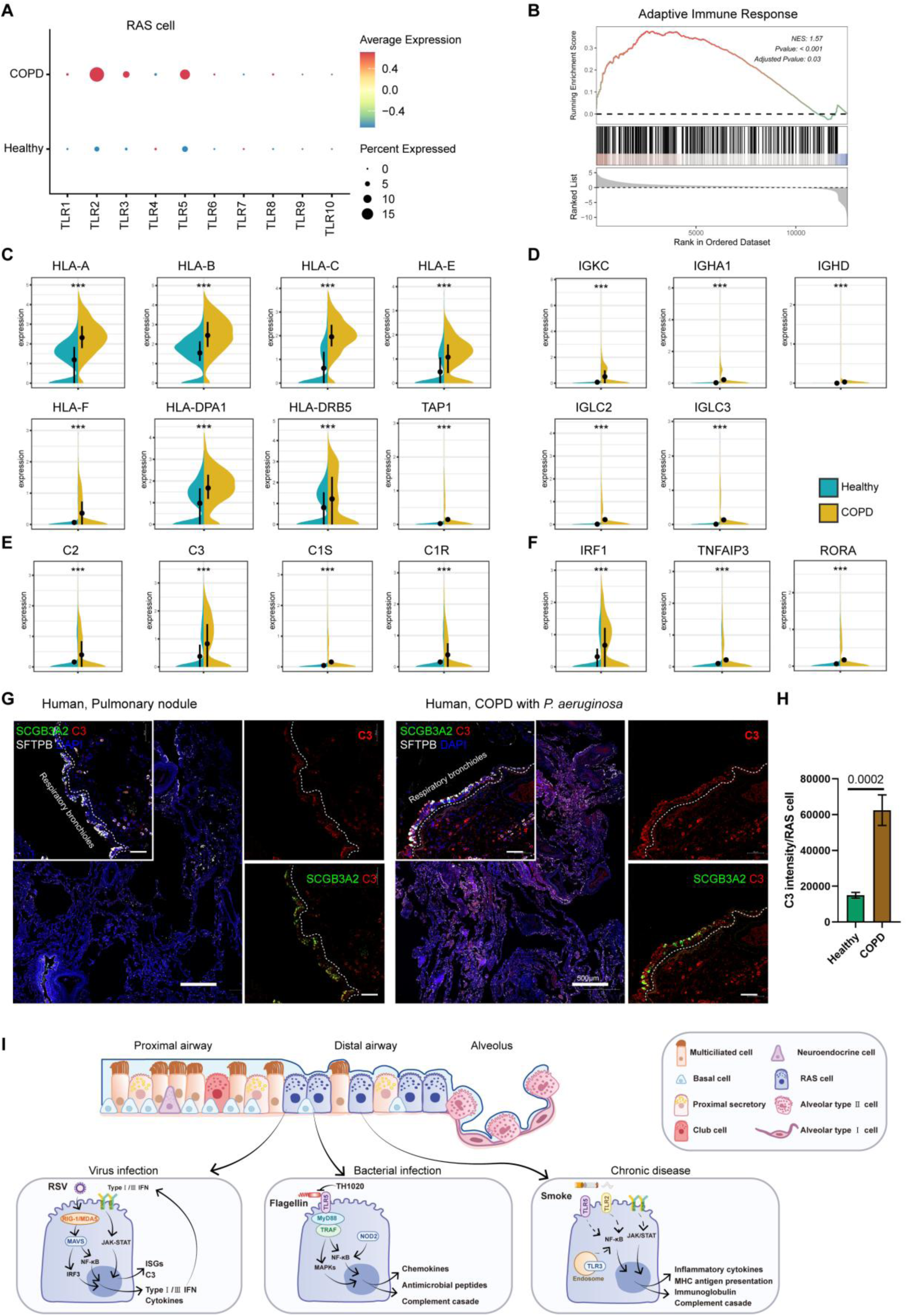
COPD is associated with enhanced immune activity in RAS cells. (**A**) Expression of Toll-like receptors (TLRs) in RAS cell clusters from COPD and healthy controls. (**B**) GSEA plot showing up-regulation of the adaptive immune response pathway in RAS cell clusters from COPD patients compared to those from healthy controls. (**C** to **F**) Violin plots illustrating expression levels of selected genes contributing to the Gene Ontology (GO) biological process ‘adaptive immune response’. (**G**) Confocal immunofluorescence images of SCGB3A2 (green), C3 (red), SFTPB (white), and DAPI (blue) in human lung sections from control individuals and *Pseudomonas aeruginosa*-infected COPD patients. White boxed areas are expanded. Scale bars, 50 μm. (**H**) Quantification of C3 fluorescence intensity per cell in healthy and COPD lung sections based on immunofluorescence analysis. Data are presented as mean ± SEM. *P*-values were calculated by unpaired two-sided Student’s *t*-test (n = 3 patients per group). (**I**) Schematic diagram summarizing the immunity role of RAS cells in acute infections (viral and bacterial) and chronic inflammatory disease (COPD).

Comparison of gene expression between healthy and COPD-derived RAS cells revealed differential enrichment of gene sets associated with extracellular matrix organization and mitochondrial ATP synthesis (fig. S11F and Supplementary table S3), suggesting that RAS cells may participate in tissue remodeling, fibrosis, and metabolic adaptation through oxidative phosphorylation. Additionally, COPD RAS cells exhibited signatures of atypical immune activation and epithelial-intrinsic immune sensing (fig. S11, G to J).

Strikingly, GSEA revealed significant enrichment of the adaptive immune response in COPD RAS cells (Fig. 6B). A broad panel of antigen presentation–related genes and MHC components was upregulated (Fig. 6C), suggesting enhanced engagement of antigen-processing and immune-interaction pathways in COPD-associated RAS cells.

Concurrently, the increased expression of immunoglobulin-related transcripts (Fig. 6D) suggests that RAS cells may contribute to local antibody transport or expression, potentially amplifying immune responses to pathogens or facilitating immune complex deposition and chronic epithelial damage. Moreover, several immune regulatory genes, including *IRF1*, *RELB*, *SOCS5*, *RORA*, and *TNFAIP3*, were elevated in COPD RAS cells (Fig. 6F), indicating a cytokine-instructed, interferon-responsive immune state. Notably, complement genes *C1R, C1S, C2*, and *C3* were markedly upregulated (Fig. 6E), with immunostaining confirming increased C3 protein expression (Fig. 6, G and H). This persistent complement activity may exacerbate epithelial injury, amplify inflammation, and sustain pathogenic immune circuits.

To determine whether these immune-associated programs were restricted to RAS cells, we compared pathway activities across major epithelial populations in the COPD dataset. Multiple epithelial populations, including RAS cells, exhibited increased immune-related pathway activity in COPD, consistent with broad epithelial immune remodeling in this chronic lung disease (Fig. S12).

Together, these results demonstrate that RAS cells undergo substantial immune-associated transcriptional remodeling in COPD and participate in the broader epithelial immune responses associated with chronic distal airway inflammation.

## Discussion

The distal respiratory airways have long eluded functional investigation due to their anatomical inaccessibility and the inability of traditional rodent models to recapitulate their biology. Here we report a robust hPSC-derived distal lung organoid system that enables developmental and functional analysis of human respiratory airway secretory cells. Building upon recent advances in hPSC-derived lung organoid systems, including the generation of expandable respiratory airway progenitors by Pezet et al.^9^ and the identification of RAS-like populations by Basil et al.^13^, our study integrates developmental lineage trajectory analysis with functional host-pathogen interaction studies within a multicellular organoid framework.

Single-cell analysis with primary human tissue from Murthy et al^14^. demonstrated that organoid-derived epithelial populations align closely with their in vivo counterparts in lineage identity, including TRB-SC, distal basal cells and neuroendocrine cells. In particular, RAS cells derived in our system share robust expression of canonical markers characteristic of primary RAS or TRB-SC populations, including SCGB3A2, SFTPB, CFTR, CEACAM6, and MUC1, supporting the preservation of the core molecular program of distal airway secretory cells. Lineage trajectory analyses further revealed a transition from RAS cells toward AT2 cells, mirroring the RAS-to-AT2 differentiation relationship previously described in primary human distal airway tissues by Basil et al.^13^.

The recent study by Pezet et al.^9^ offers an elegant hPSC-based strategy to derive expandable RAPs. In their system, SCGB3A2- and SFTPB-expressing cells emerge at the tips of branching lung organoids at later stages (∼day 75), and these cells are subsequently isolated and expanded through dissociation and passaging to generate purified RAPs populations. In contrast, our differentiation system enables the earlier emergence of RAS-like cells (∼day 35) within intact 3D organoids generating from distal lung progenitors, and thus provides a developmentally staged platform for studying the differentiation and functional investigation of RAS cells.

Despite these advances, our organoid system remains an in vitro model and does not fully capture the cellular complexity of the native lung. Certain epithelial populations, including FOXJ1⁺ ciliated cells and pre-TB-SCs, are not represented in our organoid system. In addition, a small fraction of off-target cell populations, including non–lung epithelial and neural-like cells, are present in the cultures. Recent strategies incorporating lineage purification^37^ or microengineered culture systems^38^ can generate more enriched and mature epithelial populations, representing important complementary advances in the field.

The evolutionary divergence between murine and human distal lung development significantly limits our understanding of specialized cellular niches, as the human distal lung exhibits greater branching complexity and an 8,000-fold larger surface area than its murine counterpart ^39,40^. We show that RAS cells derive from SOX9^bright^NKX2-1^bright^ bud tip progenitors and can differentiate into multiple lineages, establishing them as critical progenitors of the distal airway. Developmental lung disorders such as congenital pulmonary airway malformations (CPAM) ^41^ and bronchopulmonary dysplasia (BPD) ^42^, which involve abnormal small airway formation, may originate from disrupted differentiation trajectories of these progenitors. The plasticity of RAS cells likely evolved to enable repair and regeneration at the human distal respiratory airways ^43,44^— the initial site for gas exchange—underscoring their role in structural homeostasis.

Infections of the lower respiratory tract—ranging from respiratory bronchioles to alveoli—are leading causes of global morbidity and mortality ^45^. The unique cuboidal epithelium of the human distal airways, which lacks goblet cells and features sparse ciliation, requires alternative defense strategies distinct from proximal mucociliary clearance ^46^. Our data reveal an unexpectedly broad spectrum of immune activity in RAS cells during both acute infection and chronic lung disease (Fig. 6I). Single-cell analysis shows that RAS cells express multiple pattern-recognition receptors, including TLR2, TLR3, and TLR5, enabling them to sense diverse microbial and viral stimuli. Notably, TLR3 is a key sensor of double-stranded RNA released from viruses during viral infection ^47^. In response to RSV infection, RAS cells mount a strong interferon-driven antiviral program, and analysis of independent single-cell datasets from COVID-19 lungs reveals similar activation of antiviral and innate immune pathways in RAS-like epithelial populations. These observations support the physiological relevance of our findings and suggest that RAS cells contribute broadly to antiviral defense beyond a single pathogen context.

Increasing evidence suggests that abnormal immune responses are crucial for the initiation and progression of COPD^48^. Airway epithelial cells respond to environmental stressors and microbial signals by releasing cytokines, chemokines, and alarmins that recruit and activate immune cells^49,50^, thereby amplifying inflammatory circuits involving neutrophils and macrophages and contributing to airway remodeling and emphysema progression^51^. Complement activation represents another critical axis linking epithelial injury to immune dysregulation in COPD^52,53^. Our analyses demonstrate that RAS cells display immune activation, including enhanced adaptive immune response, complement and chemokine pathways, spanning innate and adaptive axes. These findings suggest that RAS cells participate in the broader epithelial immune responses associated with chronic airway inflammation and highlight a previously underappreciated role of distal airway epithelial cells in COPD pathobiology.

Importantly, our study identifies complement component C3 as a converging effector induced in RAS cells under both acute and chronic inflammatory conditions. While previous studies have shown that C3 hyperactivation contributes to immunopathology in viral pneumonia, including COVID-19 ^54^, murine models have demonstrated that epithelial-derived C3 is also protective during bacterial pneumonia ^55^. Our study uncovers a regulatory mechanism of C3 expression through modulation of TLR5 signaling. The TLR5–complement pathway in RAS cells represents a promising therapeutic target. Modulating TLR5 signaling or downstream complement effector functions could provide novel strategies to attenuate inflammation, preserve epithelial integrity, and restore immune balance in diseases such as viral bronchiolitis, bacterial pneumonia, and COPD.

Taken together, these findings support a role for RAS cells as dynamic and immunocompetent epithelial sentinels with prominent immune-responsive properties in the distal airway. By elucidating the developmental origin, lineage plasticity, and immune functionality of RAS cells, this study provides a framework for understanding distal airway epithelial immunity and its contribution to respiratory disease. Future research should further define how dysregulated RAS cell immunity contributes to disease progression and whether targeting these pathways may represent a potential therapeutic avenue.

## Materials and Methods

### Human Pluripotent Stem Cell

The H1 and H9 human embryonic stem cell lines (hESC) and UE005 human induced pluripotent stem cell line (hiPSC) were maintained on Matrigel-coated 6-well plates (Corning, 354277) in mTeSR1 medium (StemCell Technologies, 85850). Cells were cultured under mycoplasma-free conditions, with daily medium replacement. At ∼80% confluency (typically every 4–5 days), cells were passaged as small clusters using 0.5 mM EDTA.

### Human samples

Lung sections were obtained from the First Affiliated Hospital of Guangzhou Medical University. This study was approved by the First Affiliated Hospital of Guangzhou Medical University Institutional Review Board (ES-2023-001-02), and written informed consent was obtained from all diseased patients prior to enrollment. All patient identifying information was removed prior to use.

### Animals

Immune-deficient NOD.Cg-*Prkdc*^scid^*Il2rg*^em1^/Smoc (NSG) mice (8-10 weeks old; Shanghai Model Organisms Center, NM-NSG-001) were used in transplantation experiments. All animal procedures were approved by the Institutional Animal Care and Use Committee of Guangzhou National Laboratory (GZLAB-AUCP-2024-03-A05) and conducted in accordance with institutional guidelines.

## Method Details

### Differentiation of Lung Progenitors from hPSCs

For differentiation, human pluripotent stem cells were dissociated into single cells using Accutase (StemCell Technologies, 07920), resuspended in mTeSR1 medium supplemented with 10 μM Y27632 (MCE, HY-10071), and counted using an automated cell counter (HUAPO AP-CFcount300) with Trypan blue (Gibco; 1:1 ratio) for viability assessment. Single cells were seeded at 1 × 10⁵ cells/well on Matrigel-coated 24-well plates and maintained in mTeSR1 medium with Y27632. After 24 hours, medium was replaced with ncTarget medium (nuwacell, RP01020) without Y27632. Daily medium changes were performed until differentiation initiation at ∼70% confluency (normally 48 hours post-seeding). Definitive endoderm differentiation (day 0) followed established protocols ^56,57^. Cells were treated with 100 ng/ml activin A (MCE, HY-P70311) in MCDB131 medium (Reprocell, PM151210) and stepwise reductions of CHIR99021 (MCE, HY-10182; 3 μM on day 0, 0.3 μM on day 1, none on day 2).

After DE specification, cells were cultured for 5 days in Advanced DMEM/F12 (Gibco, 12634028) containing 1× GlutaMax (Gibco, 35050061), 10 mM HEPES (Thermo Fisher, 15630080) and Penicillin/Streptomycin (Gibco, 15070063), 1× B27 supplement (Gibco, 17504044), 1× N2 supplement (Gibco, 17502048), 500 ng/ml FGF4 (MCE, HY-P7014), 200 ng/ml noggin (MCE, HY-P7051A), 2 μM CHIR99021, 10 μM SB431542 (MCE, HY-10431) and 1 μM SAG (MCE, HY-12848). Detached anterior foregut spheroids (days 5–7) were split into two groups:

1. p-LO: Spheroids were embedded in growth factor-reduced Matrigel (Corning, 354230) and cultured in LPC medium (20 ng/ml BMP4 [MCE, HY-P7007], 10 ng/ml FGF7/KGF [MCE, HY-P70597], 10 ng/ml FGF10 [MCE, HY-P70695], 3 μM CHIR99021, 20 μM DAPT [MCE, HY-13027], and 50 nM retinoic acid [Sigma, R2625]).
2. d-LO: Spheroids were washed with PBS, dissociated into single cells with Accutase, pelleted (300 × g, 5 min), and resuspended in growth factor-reduced Matrigel. After solidification (37°C, 15 min), cultures were maintained in LPC medium with 10 μM Y27632.

Medium from both groups was refreshed every 48 hr for 7 days to induce lung progenitor differentiation.

### Airway and Alveolar Differentiation

#### Human Lung Organoid (HLO) Differentiation Medium

HLO differentiation basal medium comprised Advanced DMEM/F12 supplemented with 2.5% BSA (Proliant, 69760), 100 U/ml penicillin-streptomycin, 1× GlutaMax, 1× B27, and ITS-X (Reprocell, pb180431).

#### Airway Differentiation

Lung organoids were cultured in airway (DCIK) differentiation medium (HLO differentiation basal medium supplemented with 50 nM dexamethasone [MCE, HY-14648], 100 nM 8-bromo-cAMP [MCE, HY-12306], 100 nM 3-Isobutyl-1-methylxanthine [IBMX; MCE, HY-12318] and 10 ng/ml KGF) with medium changes every 48 hr.

#### Alveolar Differentiation

Lung organoids were cultured in alveolar (DCIK+CSB) differentiation medium (HLO differentiation basal medium supplemented with DCIK, 3 μM CHIR99021 and 10 μM SB431542) with medium changes every 48 hr.

### Histology and immunostaining

Formalin-fixed, paraffin-embedded human distal lung tissue sections (obtained from resections at the First Affiliated Hospital of Guangzhou Medical University, China) were dewaxed, rehydrated, and subjected to antigen retrieval using Improved Citrate Antigen Retrieval Solution (Beyotime, P0083) in a 95°C water bath for 15 min. Sections were washed with PBS, permeabilized and blocked using QuickBlock™ Blocking Buffer for Immunol Staining (Beyotime, P0260) for at least 15 min at room temperature (RT), followed by incubation with primary antibodies at 4 °C overnight. Sections were washed with 0.1% Triton X-100 in PBS (PBST), incubated with species-matched Alexa Fluor-conjugated secondary antibodies (1:1,000; Thermo Fisher Scientific) for 2 hr at RT, counterstained with Hoechst 33342 (1:1,000, Thermo Scientific, 62249), and mounted with fluorescence mounting medium (DAKO, S3023). Images were acquired using a Zeiss LSM980 confocal microscope.

For whole-mount staining, organoids were extracted from Matrigel using Cell Recovery Solution (Corning, 354253), fixed in 4% paraformaldehyde (PFA) for 2 hr at 4°C, permeabilized, and stained as above. Primary antibodies used were rabbit anti-NKX2-1 (1:300, Abcam, ab76013), goat anti-SOX2 (1:300, Abcam, ab239218), mouse anti-SOX9 (1:500, Abcam, ab76997), mouse anti-Ac-Tub (1:1000, Sigma-Aldrich, T7451), rabbit anti-TP63 (1:200, Abcam, ab124762), mouse anti-VIM (1:100, Santa Cruz Biotechnology, sc-6260), mouse anti-SFTPB (1:200, Santa Cruz Biotechnology, sc-133143), goat anti-SCGB3A2 (1:500, R&D Systems, AF3545), mouse anti-MUC5AC (1:500, Abcam, ab3649), rabbit anti-SFTPC (1:200, Seven Hills Bioreagents, WRAB-9337), goat anti-AGER (1:200, R&D Systems, AF1145), rabbit anti-TLR5 (1:500, abcam, ab300038), mouse anti-HUNU (1:300, Abcam, ab191181), rabbit anti-CXCL10 (1:100, Invitrogen, MA532674), mouse anti-CXCL8 (1:500, Invitrogen, M801), rabbit anti-RSV-F (1:200, Sino Biological, 11049-R302). Organoids were incubated with Alexa Fluor 647 phalloidin (1:1000, Thermo Fisher Scientific, A22287), with the corresponding secondary antibodies Alexa Fluor 488 Donkey anti-Rabbit (Thermo Fisher Scientific, A21206), Alexa Fluor 546 Donkey anti-Rabbit (Thermo Fisher Scientific, A10040), Alexa Fluor 488 Donkey anti-Mouse (Thermo Fisher Scientific, A21202), Alexa Fluor 546 Donkey anti-Mouse (Thermo Fisher Scientific, A10036), Alexa Fluor 405 Donkey anti-Goat (Thermo Fisher Scientific, A48259), Alexa Fluor 647 Donkey anti-Goat (Thermo Fisher Scientific, A21447) in blocking buffer containing Hoechst 33342 solution (20 mM) (1:1,000). Organoids were transferred to a confocal dish, and images were acquired using the Zeiss LSM980 confocal microscope.

### RSV Infection of Organoids

RSV strain Long was propagated as described ^58,59^. RSV infection was performed in D40 d-ALOs. d-ALOs were infected at a multiplicity of infection (MOI) of 0.1 for 2 hours at 37°C. Organoids were washed three times with DPBS (Hyclone, AH30017338) and cultured in fresh alveolar medium for 24–72 hours. Organoids and supernatants were collected at designated time points for downstream analysis.

### Assessment of Cytokine Production

To evaluate cytokine secretion, FLA-PA stimulation was performed in D40 d-ALOs. d-ALOs were treated with varying concentrations of FLA-PA (InvivoGen, tlrl-pafla). Supernatants were collected at designated timepoints, aliquoted, and stored at −80°C until batch analysis. Cytokine levels were quantified using a magnetic Luminex® multiplex assay (R&D Systems, LXSAHM-08) according to the manufacturer’s instructions. Briefly, serial dilutions of cytokine standards were prepared. Antibody magnetic bead mix was dispensed into a 96-well plate, followed by the addition of standards and experimental samples. The plate was incubated overnight at 4°C with shaking. After three washes, detection antibodies were introduced and incubated at RT for 30 minutes with shaking. The plate was washed three times. Streptavidin-PE was subsequently added and incubated under identical conditions. Following a final wash cycle, 100 μl of wash buffer was dispensed into each well. Fluorescence intensity was measured using a Luminex X-200 system, and acquired data were processed with Milliplex Analyst software (v5.1).

### Mouse Kidney Capsule Transplantation

All animal procedures were approved by the Institutional Animal Care and Use Committee of Guangzhou National Laboratory and conducted in accordance with institutional guidelines. NOD.Cg-*Prkdc*^scid^*Il2rg*^em1^/Smoc (NSG) mice (8-10 weeks old; Shanghai Model Organisms Center, NM-NSG-001) were housed under specific pathogen-free conditions. d-ALO cells mixed with 5 μL Matrigel were implanted beneath the right kidney capsule under anesthesia. At 4 weeks post-transplantation, grafts were excised, fixed in 4% PFA overnight, dehydrated in 30% sucrose overnight, and embedded in Optimal Cutting Temperature (OCT) compound or paraffin for immunofluorescence or hematoxylin and eosin (H&E) staining, respectively.

For bacterial infection, PAO1 (MiaoLingBio, T0133) was cultured overnight in LB medium at 37°C with shaking. The bacterial suspension was centrifuged, washed three times with PBS, and resuspended in sterile saline to an optical density OD600 of 1.0, corresponding to 1 × 10^9^ colony-forming units (CFU) per mL. For experimental use, the suspension was diluted to 1 × 10^6^ CFU/mL, pelleted by centrifugation (10,000 × g, 5 min, 4°C), and subjected to four freeze-thaw cycles to achieve bacterial lysis. At 4 weeks post-transplantation, mice were anesthetized and a posterior left incision was made to expose the engrafted d-ALO. A 50 µL aliquot of PAO1 lysate was injected into the d-ALO lumen using a sterile 0.5-mL insulin syringe. The d-ALO was repositioned into the peritoneal cavity, and the incision was closed via a two-layer suturing technique. For pain control, mice were given an injection with carprofen subcutaneously (4 mg/kg, Zoetis). At 48 hours post-injection, mice were euthanized. Engrafted d-ALO tissues and adjacent kidney samples were harvested for subsequent analysis.

### RNA Isolation and qPCR

Total RNA was extracted using the RNeasy Kit (Qiagen, 74106) following the manufacturer’s instructions. qPCR was performed using Bio-Rad CFX384 Real-Time PCR Detection System, with primers listed in Supplementary Table 1. Biological and technical replicates were analyzed as previously described ^60^.

### Flow Cytometry

Lung organoids were dissociated into single cells using TrypLE (Gibco, 12605028) at 37°C. For AT2 sorting, cells were stained 30 mins with primary antibody (mouse anti-HTⅡ-280 [1:150, TerraceBiotech, TB-27AHT2-280]) at 4°C, followed by secondary antibody incubation (45 minutes, RT). Cells were sorted using Sony MA900 Cell Sorter. For NKX2-1 assessment, after dissociation, cells were fixed in 1% PFA for 15 minutes at 4°C, and permeabilized with 1× Perm/Wash Buffer (BD Biosciences, 554723) for 20 minutes at RT. Cells were stained under identical condition using anti-NKX2-1 antibody (1:300, Abcam, ab76013) and analyzed by Beckman CytoFLEX S flow cytometer. Data were analyzed using FlowJo v10.6.2. or CytExpert.

### Forskolin-Induced Organoid Swelling

d-AWOs at day 35–45 were used for swelling assays. Prior to treatment, organoids were passaged into 3 μL growth factor-reduced Matrigel droplets and cultured in the absence of 8-bromo-cAMP and IBMX for at least one day. For swelling analysis, organoids were incubated with 10 mM forskolin (Aladdin Scientific, F127328-500mg) in fresh medium for 5–24 hours at 37°C under 5% CO₂. Brightfield images of entire droplets were captured at 0, 5, 7, and 24 hours post-treatment using a High-Content Imaging System (PerkinElmer, Operetta CLS) and stitched with Harmony software. Swelling area ratios (post-treatment area / baseline area) were quantified in ImageJ from replicate wells, with the initial area (t = 0) normalized to 1. Statistical analyses included 20 organoids per experimental replicate.

### Transmission Electron microscopy

d-ALOs were fixed in 2.5% glutaraldehyde for 30 minutes at RT, washed with 0.1 M phosphate buffer (pH 7.4), and postfixed in 1% osmium tetroxide for 1.5 hours at RT. After repeated phosphate buffer washes, samples were dehydrated in a graded ethanol series (50%, 70%, 80%, 90%, 100%) at RT. Tissues were infiltrated with propylene oxide, followed by a 1:1 propylene oxide/Epon mixture (2–4 hours, RT), then pure Epon resin (1 hour, RT). Samples were embedded in Epon resin, polymerized at 40°C (24 hours) and 60°C (12 hours), and sectioned (70–90 nm). Ultrathin sections were stained with uranyl acetate and lead citrate and imaged on a FEI Tecnai Transmission Electron Microscope.

### Live-Cell Imaging of Lamellar Bodies

d-ALOs were incubated with β-BODIPY FL C12-HPC (1 μM; Life Technologies, D3792) in alveolar differentiation medium for 24 hours at 37°C/5% CO₂. Organoids were washed and stained with LysoTracker Red DND-99 (100 nM; Life Technologies, L7528) for 30 min at RT, followed by two washes. Live imaging was performed at 37°C using a Zeiss LSM980 confocal microscope.

### Organoid scRNA-seq

#### Sample Preparation and Library Construction

Single-cell suspensions were generated from lung organoids, and viability was assessed using a Luna Dual Fluorescence Cell Counter. Libraries were prepared with the Chromium Single Cell 3′ Library and Gel Bead Kit v3 (10x Genomics) following the manufacturer’s protocol. Briefly, single-cell suspensions were loaded into a Chromium Controller to produce Gel Bead-in-Emulsions (GEMs). Following reverse transcription, cDNA amplification, and library construction, sequencing was performed on an Illumina NovaSeq 6000 platform (PE150; Annoroad Gene Technology).

#### Data Preprocessing

Raw sequencing data were processed via the Cell Ranger pipeline (v4.0.0). Reads were aligned to the GRCh38 human genome, and gene expression matrices were generated. Low-quality cells (fewer than 500 detected genes or >10% mitochondrial genes) were excluded. Doublets were identified and removed using DoubletFinder (v2.0.4).

#### Data Analysis with Seurat

The filtered gene expression matrix was analyzed using the Seurat (v5.1.0). After quality control, the datasets were merged independently using the ‘Seurat::merge()’ function. Data normalization was performed using the “NormalizeData” function with a scale factor of 10,000. Highly variable genes were identified using the “FindVariableFeatures” function, and the data were scaled using the “ScaleData” function. Principal Component Analysis (PCA) was conducted to reduce dimensionality. Batch effects were then corrected by integrating the data using the “Harmony” method. Cells were clustered using the “FindNeighbors” and “FindClusters” functions. The clusters were visualized using Uniform Manifold Approximation and Projection (UMAP) with the “RunUMAP” function. Marker genes for each cluster were identified using the “FindAllMarkers” function with a log-fold change threshold of 0.25.

#### Cell-Type Annotation

Cell-type assignment was performed using a hierarchical annotation strategy to precisely resolve cellular heterogeneity within the organoids. Initially, major cellular compartments were defined by manual evaluation of canonical lineage-defining marker genes. This broad classification allowed for the identification and distinction of the definitive epithelial lineage from non-epithelial populations present in the organoid system. To validate these broad annotations, we performed differential expression analysis using the ‘FindAllMarkers’ function in Seurat (Wilcoxon Rank Sum test) to confirm that the top upregulated genes for each cluster corresponded to their respective lineage identities. To achieve high-resolution characterization of the lung epithelium, the epithelial compartment was computationally subsetted from the global dataset. This subsetted object underwent a second round of data processing, including variable feature selection, scaling, Principal Component Analysis (PCA), and UMAP dimensionality reduction. The resulting epithelial subclusters were then annotated using a comprehensive strategy: first, cluster-specific differentially expressed genes (DEGs) were identified using the ‘FindAllMarkers’ function; and second, these signatures were cross-referenced with known stage-specific and cell-type-specific marker genes to assign definitive biological identities.

#### Differential Expression and Pathway Analysis

Differentially expressed genes (DEGs) between clusters were identified using the “FindMarkers” function, calculating fold change and performing Wilcoxon Rank Sum tests. Gene Ontology (GO) and pathway enrichment analyses were performed using clusterProfiler (v4.14.4) to interpret the biological significance of the identified markers.

#### Trajectory Inference Analysis

Trajectory analysis was conducted using Monocle3 (v1.3.7). The UMAP coordinates obtained from Seurat were utilized as the input for Monocle3 to construct the trajectory. Cells were ordered along the pseudotime trajectory based on the learned principal graph, allowing for the identification of dynamic gene expression changes across different stages of the trajectory.

#### Single-cell reference mapping

Reference-based mapping was performed using the Seurat R package. The dataset from Sun et al. was used as the query dataset, and the dataset from Murthy et al. (2022) served as the reference atlas. The query dataset was preprocessed using the standard Seurat workflow, including normalization, identification of highly variable genes, data scaling, and principal component analysis (PCA). Transfer anchors between the reference and query datasets were identified using the FindTransferAnchors function based on the PCA reduction of the reference dataset. Reference cell-type annotations were then transferred to the query cells using the MapQuery function, which performs label transfer, embedding integration, and projection of query cells into the reference UMAP space. Briefly, MapQuery internally executes TransferData to assign predicted cell-type labels, IntegrateEmbeddings to align the query cells with the reference embedding, and ProjectUMAP to project query cells onto the precomputed reference UMAP (refUMAP) for visualization. Cell-type identities of query cells were assigned based on the highest prediction scores obtained during label transfer.

#### kNN-based annotation overlap analysis

To quantify transcriptomic similarity between two datasets, a k-nearest neighbor (kNN)–based annotation overlap analysis was performed. Cells from both datasets were embedded into a shared low-dimensional space using the CCA-derived UMAP reduction generated during Seurat integration. A kNN graph was constructed using the FindNeighbors function in Seurat with k = 30 based on the CCA-derived embedding. For each query cell, its nearest neighbors were identified within the combined dataset. To evaluate cross-dataset similarity, only neighbors belonging to the reference dataset were retained for downstream analysis. For each query cell type, the reference neighbors of all corresponding query cells were aggregated, and the distribution of reference cell-type annotations among these neighbors was calculated. The resulting proportions reflect the degree of transcriptomic similarity between query and reference cell populations. The overlap matrix (query cell types versus reference cell types) was visualized as a heatmap using the pheatmap R package, where higher values indicate stronger similarity between the corresponding cell populations.

### Statistical analysis

Statistical tests (one-way ANOVA, two-way ANOVA, unpaired Student’s *t*-test) were performed in GraphPad Prism (v8.0.2). Data represent biologically independent replicates (≥3 experiments, except scRNA-seq). Results are expressed as mean ± SEM. All tests were two-sided (*α* = 0.05).

## Acknowledgments

We thank the Imaging and Cell Technology Platforms, Guangzhou Laboratory for the confocal and flow cytometry. This work was supported by the National Natural Science Foundation of China (82171419 to J.S.), Major Project of Guangzhou National Laboratory (MP-GZNL2025C03007 to H.L.), National Key Research and Development Program of China (2021YFA1101304 to H.L.), National Key Research and Development Program of China (2020YFA0908200 to H.L.).

## Author contributions

Conceptualization: J.S., H.L.

Formal analysis: J.S., H.S., S.J.

Investigation: J.S., H.S., X.X.

Visualization: J.S., H.S., S.J.

Funding acquisition: H.L., J.S.

Project administration: J.S., H.L.

Resources: D.W., W.G., H.Y., H.Z., H.M., J.D.

Supervision: H.L., T.X., J.Z., W.K.Z.

Writing – original draft: J.S.

Writing – review & editing: H.L., W.K.Z., J.S.

## Declaration of interests

The authors declare no competing interests.

## Data and code availability

Organoids scRNA-seq data that support the findings of this study were deposited in the Gene Expression Omnibus under accession code GSE334709, and are publicly available.

## Supplementary Figure

**Fig. S1. Characterization of proximal and distal lung organoid differentiation. A**, Differentiation protocol workflow for generating proximal lung organoids (p-LOs) from hPSCs. **B**, Bright-field images comparing organoid morphology under untreated versus dissociation conditions during lung progenitor cell (LPC) induction. Scale bars, 500 µm (far left and middle two panels), 200 µm (right panels). Representative of three biologically independent experiments. **C**, Phase-contrast images of day 8 p-LOs and d-LOs counterstained with Hoechst. Scale bars, 500 µm. **D**, Organoid yield quantification (number per Matrigel droplet). **E**, Organoid diameter measurements. **F**, Representative flow cytometry analysis of NKX2-1 expression in p-LOs versus d-LOs. n = 3 independent experiments. **G**–**H**, Percentage (**G**) and mean fluorescence intensity (MFI) (**H**) of NKX2-1^+^ cells. **I**, qPCR analysis of lung-specific and endodermal lineage markers in p-LOs and d-LOs derived from hESC lines (H1, H9) and induced pluripotent stem cells (hiPSCs). **J**, Integrated UMAP projection of p-LOs and d-LOs colored by annotated major cell types (left) and experimental conditions (right). **K**, Dot plot displaying canonical lineage markers for epithelial, mesenchymal, and neural cell populations. **L**, Proportional distribution of major cell types in d-LOs versus p-LOs. **M**, Epithelial subcluster marker expression distinguishing d-LPC, p-LPC, neuroendocrine cells, cycling cells, and hindgut-like cells. **N-O**, Gene Ontology (GO) enrichment analysis of biological processes enriched in p-LPC (**N**) and d-LPC (**O**) clusters. **P**, Expression patterns of proximal and distal lineage markers from fetal lung ^15^ in p-LPC and d-LPC clusters. **Q**–**S**, Heatmaps displaying the overlap coefficients of gene expression signatures from p-LPC and d-LPC clusters compared with cell type annotations from published human fetal lung scRNA-seq datasets by Cao et al., ref. 15 (**Q**), Sountoulidis et al., ref. 24 (**R**), and He et al., ref. 23 (**S**). **D**,**E**,**G**,**H**,**I**, Data are presented as mean ± SEM (n=3 biological replicates). *P*-values calculated using two-tailed Student’s *t*-test.

**Fig. S2. d-LOs generate distal airway and alveolar lineages containing RAS cells. A**, Differentiation timeline for generating airway and alveolar organoids from proximal (p-LO) and distal (d-LO) lung organoids. Arrows indicate time points for qPCR and scRNA-seq analyses. **B**, qPCR quantification of lung lineage markers in p-LO- and d-LO-derived organoids. Lung epithelial marker: *NKX2-1*; Airway markers: *SCGB3A2*, *TP63*, *MUC5AC*, *SCGB1A1*, *FOXJ1*; alveolar markers: *SFTPB*, *AGER*, *PDPN*, *SFTPC*, *LAMP3*. Data are presented as mean ± SEM (n=3). *P*-values calculated by one-way ANOVA with Tukey’s multiple comparison test. **C**, Immunofluorescence co-staining of lineage markers in organoids: basal (TP63), distal secretory (SCGB3A2, SFTPB), multiciliated (Ac-Tub), mesenchymal (VIM), goblet (MUC5AC), AT2 (SFTPB, SFTPC, NKX2-1), and AT1 (AGER). Insets on select images highlight protein expression in specific cells. Scale bars, 50 µm (low-magnification images), 20 µm (expanded insets). Representative of three independent experiments. **D**, Schematic of regional lineage specification from proximal airway and distal airway/alveolar organoids. **E-F**, UMAP projections of single-cell RNA sequencing data from d-LOs and organoids under differentiation conditions, colored by all (**E**) or individual differentiation time/media conditions (**F**). **G**, Annotated UMAP distinguishing major cell types across differentiation conditions. **H**, Dot plot of canonical markers defining cell clusters in (**G**). **I**, Stacked bar plot quantifying the relative proportions of annotated cell types across different differentiation conditions.

**Fig. S3. Functional validation of distal lung organoid derivatives and cross-dataset comparison. A**, Time-lapse phase-contrast imaging of forskolin-induced swelling in distal airway organoids (d-AWO) derived from d-LOs. Scale bars, 1 mm (upper panel), 500 µm (lower panel). **B**, Quantification of normalized luminal swelling area in d-AWOs over 24 hours. Bars represent mean ± SEM (n=20 organoids). *P*-values calculated using two-tailed Student’s *t*-test. **C**, Schematic of apical-out (AO) polarity model (left) and representative microscopy showing cilia in AO-d-AWO. Scale bar, 10 µm. **D**, Transmission electron microscopy (TEM) of d-ALOs showing lamellar body-like structures. Scale bar, 500 nm. **E-F**, BODIPY-labeled phosphatidylcholine uptake in Matrigel-embedded (**E**) and apical-out (**F**) d-ALOs. Scale bars, 20 µm. Representative of three independent experiments. **G**, Reference mapping of Sun et al. scRNA-seq data onto the Murthy et al. (ref. 14) human fetal lung atlas using a query-to-reference projection. Left panel: cells colored by predicted cell types from Murthy et al.; Right panel: cells colored by Sun et al. original annotations. **H**, Stacked bar plot quantifying the composition of predicted Murthy et al. cell types within each annotated cluster from Sun et al. **I**–**J**, Integrated UMAP projection of combined datasets (**I**) and split views (**J**) showing the alignment of Sun et al. and Murthy et al. cells in a shared embedding space. **K**, Heatmap displaying kNN-based annotation overlap coefficients between Sun et al. clusters (columns) and Murthy et al. cell types (rows).

**Fig. S4. Lineage potential of RAS cells in d-LO-derived organoids. A-B**, UMAP projections of single-cell transcriptomes from d-LOs and their derivatives. Cells colored by experimental conditions (**A**) or annotated cell types (**B**). **C**, Proportional distribution of cell types in d-LOs versus d-LO-derived organoids. **D**, Schematic showing trajectory analysis identifying four differentiation paths from RAS cells to PNEC, basal, and AT2 lineages as well as from p-LPC to ciliated cell. **E**-**H**, UMAP projections depicting four cellular trajectories with color indicating progression (top) and heatmaps of trajectory-defining gene expression (bottom) across pseudotime for each lineage relationship.

**Fig. S5. Independent validation of lineage trajectory using Slingshot and diffusion pseudotime analyses. A**, UMAP projection of integrated d-LO and d-LO-derived organoid epithelial cells with Slingshot-inferred lineage trajectories overlaid. Cell types are annotated by color. **B**, Pseudotemporal ordering of epithelial subtypes along the Slingshot-derived trajectory. Cells are colored by pseudotime progression. **C**-**F**, UMAP projections depicting four distinct differentiation trajectories with color indicating progression for each lineage relationship. **G**, Diffusion pseudotime (DPT) analysis of integrated epithelial populations. Cells colored according to diffusion pseudotime values, showing the global trajectory progression. **H**, The same DPT embedding colored by annotated cell types. Arrows indicate the predicted developmental progression.

**Fig. S6. Enrichment of immune- and host defense–associated pathways across epithelial populations in organoids. A-K**, Violin plots showing module scores for selected immune-and host defense–related pathways across major epithelial cell populations. Pathways analyzed include positive regulation of pattern-recognition receptor signaling (**A**), positive regulation of cytokine production (**B**), complement activation (**C**), cellular response to interferon-γ (**D**), positive regulation of inflammatory response (**E**), regulation of viral entry into host cells (**F**), regulation of viral processes (**G**), antibacterial humoral response (**H**), defense response to bacterium (**I**), positive regulation of interleukin-2 production (**J**), and antimicrobial humoral response (**K**). RAS cells showed prominent enrichment of multiple immune-sensing, inflammatory, and host defense programs.

**Fig. S7. Immune defense capacity of RAS cells in d-LO derived organoids infected by RSV. A**, qPCR analysis of *CXCL8*, *CXCL10*, *IL-6*, and *TNF-α* expression in d-ALOs infected with RSV (MOI 0.1) at 24 and72 hours post-infection (hpi). Data are presented as mean ± SEM (n=3). *P*-values calculated using two-tailed Student’s *t*-test. **B**, Integrated UMAP projection of scRNA-seq data from mock- and RSV-infected organoids, colored by annotated cell types (left) and infection status (right). **C**, Gene Ontology (GO) enrichment analysis of shared DE genes, highlighting “defense response to virus” as the top pathway. The full name of the term marked by an asterisk is ‘Antigen Presentation: Folding, assembly and peptide loading of class I MHC’. **D**, Heatmap of 126 common DE genes across clusters in RSV-infected d-ALOs. **E**–**G**, Cell cycle analysis of scRNA-seq data from mock- and RSV-infected organoids. UMAP projection colored by cell cycle phase (**E**), stacked bar plot quantifying the proportion of cells in G1, G2M, and S phases (**F**), and heatmap of cell cycle regulatory genes (**G**) in RAS^high^ cell, RAS^low^ cell, and RAS-mock groups. **H**, Dot plot showing the expression of canonical lineage markers identifying distinct cell populations in the single-cell lung atlas of lethal COVID-19 (Melms et al., ref. 32). **I**, Heatmap validating the expression of the 121 core antiviral signature genes in airway club cells from control and COVID-19 patients (Melms et al., ref. 32).

**Fig. S8. Cell-type–resolved analysis of RSV infection responses across epithelial populations. A**, UMAP projection of RSV-infected cells colored by the 126 RSV-response gene signature score. **B**, Violin plot comparing RSV-response signature scores among RAS cells, PNECs, and non-epithelial populations. **C**, Reference-based mapping of EpCAM⁺ cells onto the integrated distal lung epithelial atlas. **D**, Cell-type composition of mock and RSV-infected organoid cultures. **E**–**G**, Distribution of RSV transcripts across epithelial populations. UMAP projection colored by RSV transcript abundance (**E**), identification of RAS Cell^high^ and RAS Cell^low^ subsets (**F**), and dot plot showing expression of RSV genes across epithelial cell types (**G**). **H**. UMAP projection colored by the 126 RSV-response signature score. **I**. Comparison of signature scores between mock and RSV-infected conditions across epithelial populations. **J**. Comparison of RSV-response signature scores among epithelial cell types in infected cultures.

**Fig. S9. scRNA-seq analysis of RAS cells upon TLR5 activation. A**, Dot plot showing the expression of Toll-like receptors (TLRs) across all cell clusters (Fig. S2H). **B**, Violin plots comparing TLR5 expression in RAS cells under indicated experimental conditions. **C**–**D**, Validation of cell type markers and TLR5 expression in the human fetal lung atlas from Quach et al. (ref. 34). **C**, Dot plot of lineage markers identifying distinct cell populations; the red box highlights the gene signature of SCGB3A2^+^SFTPB^+^CFTR^+^ cells. **D**, Dot plot displaying TLR5 expression levels across the cell types defined in (**C**). **E**, Cytokine/chemokine secretion (CXCL8, CXCL10, IL-6, IL-1β, TNF-α) from d-ALOs stimulated with increasing concentrations of FLA-PA protein (1–50 µg/ml) over 6–72 hours. Data are presented as mean ± SEM (n=3 biological replicates). *P*-values calculated using two-way ANOVA. **F**, UMAP projections of single-cell transcriptomes from control, FLA-PA-treated, and FLA-PA+TH1020-treated d-ALOs. Cells colored by experimental conditions (right) or annotated cell types (left). **G**, Proportional distribution of cell types in control, FLA-PA-treated, and FLA-PA+TH1020-treated d-ALOs. **H**, Cluster defining genes within (**F**) are shown in dot plot format. **I**, Heatmap from pseudo-bulk analysis of scRNA-seq data showing changes in mRNA expression of TLR signaling pathway related genes in RAS cells stimulated with FLA-PA (1μg/ml) with or without TH1020 (1μM) for 72h. **J**–**L**, Cell cycle analysis of the cell populations shown in (**F**). UMAP colored by cell cycle phase (**J**), stacked bar plot quantifying phase proportions (**K**) and heatmap of cell cycle regulatory genes (**L**) in RAS cell cluster across treatment groups.

**Fig. S10. Specificity of FLA-PA-induced immune responses in RAS cells. A**-**F**, GSEA reveals significantly upregulated items in RAS cells after stimulated with FLA-PA, indicating that these cells activated a series of functions related to immune recognition (**A**), leukocyte recruitment (**B**), antibacterial response (**C**), cytokine and receptor signal regulation (**D**), cytoskeleton remodeling and secretion regulation (**E**), and programmed cell death and immune regulation (**F**). **G**, Flow cytometry gating strategy for AT2 cell isolation using HTII-280 surface marker. **H**, qPCR validation of *TLR5*, *SFTPC*, *SFTPB*, and *SCGB3A2* expression in sorted HTII-280+ AT2 cells, confirming high alveolar (SFTPC) and low immune (TLR5) signatures. **I**, Comparative RT-PCR analysis of *CXCL8* and *IL-6* in whole d-ALOs versus sorted HTII-280+ AT2-enriched organoids after 72 h FLA-PA stimulation (1 µg/ml). **J**, GSEA plots evaluating “Regulation of Toll Like Receptor Signaling Pathway” (top row) and “Humoral Immune Response” (bottom row) signatures across other cell types (Basal, Ciliated, PNEC, Cycling, non-lung epithelial, Mesenchymal). **H**,**I**, Data are presented as mean ± SEM (n=3). *P*-values calculated using two-tailed Student’s *t*-test.

**Fig. S11. scRNA-seq analysis of RAS cells from COPD. A**, Reanalysis of scRNAseq dataset from normal and COPD peripheral samples (Basil *et al*., ref. 13) and subset of epithelium showing expected epithelial populations. The COPD cohort consists of patients with severe COPD as defined by FEV1 values from the source study (Basil et al., ref. 13), including one patient with GOLD stage III and three patients with GOLD stage IV COPD. **B**, Cluster defining genes within the epithelium are shown in dot plot format. **C**, Stacked bar graphs showing the normal and COPD patients contribution to each epithelial cell cluster. **D**, Heatmap from pseudo-bulk analysis of scRNA-seq data showing significant changes in mRNA expression of TLR signaling pathway related genes in RAS cells from normal and COPD donors. **E**, Immunofluorescence of SCGB3A2 (green), CXCL8/CXCL10 (red), SFTPB/TLR5 (white), and DAPI (blue) in human lung sections from control and *Pseudomonas aeruginosa*-infected COPD patients. White boxed area is expanded. Scale bars, 50 μm. **F**, Top 15 GO terms enriched in DEGs between RAS cells from COPD versus from healthy controls. Significantly enriched terms highlight increased mitochondrial ATP synthesis, extracellular matrix organization, and immune-related processes, suggesting functional adaptation of RAS cells under chronic disease conditions. **G**-**J**, GSEA reveals significantly upregulated immune related items in RAS cells from COPD, including antibody-related immune function (**G**), antigen recognition (**H**), Cytokine signaling and immune regulation (**I**), and endocytosis (**J**).

**Fig. S12. Comparison of immune-associated pathway activities across epithelial populations in healthy and COPD lungs. A**-**E**, Violin plots showing gene signature enrichment scores for representative immune-related pathways across major epithelial cell populations from healthy donor and COPD lungs. Pathways analyzed include adaptive immune response (**A**), complement activation (**B**), cytokine activity (**C**), positive regulation of receptor-mediated endocytosis (**D**), and antigen binding (**E**).

## Notes

### Competing Interest Statement

The authors have declared no competing interest.

### Summary of Updates

We have added several new figures and corrected the GEO accession number to GSE334709.

