## supplementary_materials for "hPSC-Derived Distal Lung Organoids Reveal Respiratory Airway Secretory Cells Acting as Immune Sentinels in Human Distal Airways"

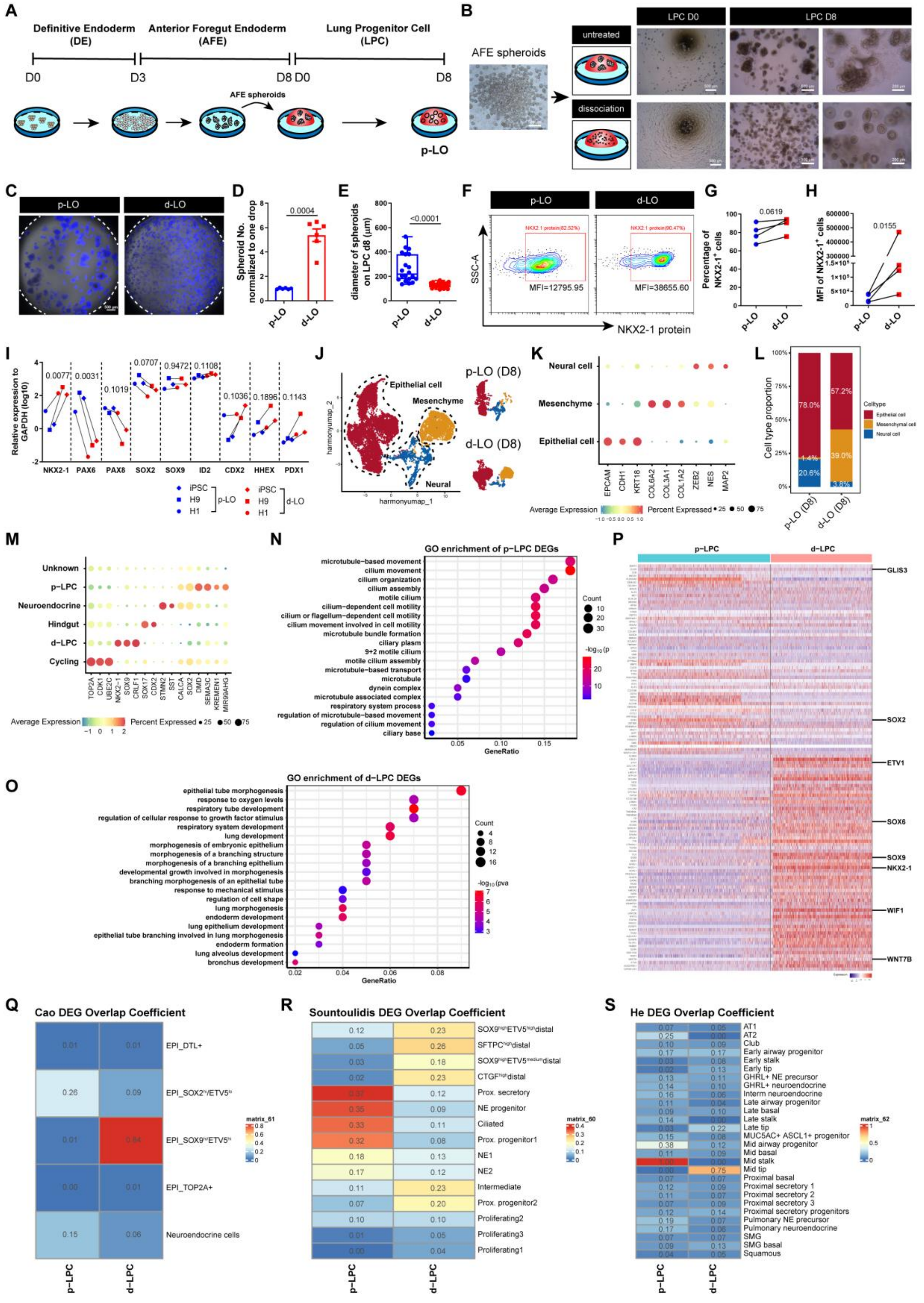
A
Definitive Endoderm (DE)
Anterior Foregut Endoderm (AFE)
Lung Progenitor Cell (LPC)
D0
D3
D8 D0
D8
AFE spheroids
p-LO
B
AFE spheroids
untreated
dissociation
LPC D0
LPC D8
C
p-LO
d-LO
D
Spheroid No. normalized to one drop
0.0004
E
diameter of spheroids on LPC d8 (μm)
<0.0001
F
p-LO
d-LO
SSC-A
NKX2-1 protein
MFI=12795.95
MFI=38655.60
G
Percentage of NKX2-1+ cells
0.0619
H
MFI of NKX2-1+ cells
0.0155
I
Relative expression to GAPDH (log10)
NKX2-1 PAX6 PAX8 SOX2 SOX9 ID2 CDX2 HHEX PDX1
iPSC H9 H1 p-LO
iPSC H9 H1 d-LO
J
Epithelial cell
Mesenchyme
Neural
harmonyumap_1
harmonyumap_2
p-LO (D8)
d-LO (D8)
K
Neural cell
Mesenchyme
Epithelial cell
EPCAM CDH1 KRT18 COL6A2 COL3A1 COL1A2 ZEB2 NES MAP2
Average Expression
Percent Expressed
L
Cell type proportion
78.0%
57.2%
39.0%
20.6%
Celltype
Epithelial cell
Mesenchymal cell
Neural cell
p-LO (D8)
d-LO (D8)
M
Unknown
p-LPC
Neuroendocrine
Hindgut
d-LPC
Cycling
TOP2A CDK1 UBE2C NKX2-1 SOX9 CRLF1 SOX17 CDX2 STMN2 SST CALCA SOX2 DMD SEMA3C KREMEN1 MIR99AHG
Average Expression
Percent Expressed
N
GO enrichment of p-LPC DEGs
microtubule-based movement
cilium movement
cilium organization
cilium assembly
motile cilium
cilium-dependent cell motility
cilium or flagellum-dependent cell motility
cilium movement involved in cell motility
microtubule bundle formation
ciliary plasm
9+2 motile cilium
motile cilium assembly
microtubule-based transport
microtubule
dynein complex
microtubule associated complex
respiratory system process
regulation of microtubule-based movement
regulation of cilium movement
ciliary base
GeneRatio
Count
P
p-LPC
d-LPC
GLIS3
SOX2
ETV1
SOX6
SOX9
NKX2-1
WIF1
WNT7B
O
GO enrichment of d-LPC DEGs
epithelial tube morphogenesis
response to oxygen levels
respiratory tube development
regulation of cellular response to growth factor stimulus
respiratory system development
lung development
morphogenesis of embryonic epithelium
morphogenesis of a branching structure
morphogenesis of a branching epithelium
developmental growth involved in morphogenesis
branching morphogenesis of an epithelial tube
response to mechanical stimulus
regulation of cell shape
lung morphogenesis
endoderm development
lung epithelium development
epithelial tube branching involved in lung morphogenesis
endoderm formation
lung alveolus development
bronchus development
GeneRatio
Count
Q
Cao DEG Overlap Coefficient
EPI_DTL+
EPI_SOX2hi/ETV5lo
EPI_SOX9hi/ETV5hi
EPI_TOP2A+
Neuroendocrine cells
matrix_61
p-LPC
d-LPC
R
Sountoulidis DEG Overlap Coefficient
SOX9highETV5highdistal
SFTPChighdistal
SOX9highETV5mediumdistal
CTGFhighdistal
Prox. secretory
NE progenitor
Ciliated
Prox. progenitor1
NE1
NE2
Intermediate
Prox. progenitor2
Proliferating2
Proliferating3
Proliferating1
matrix_60
p-LPC
d-LPC
S
He DEG Overlap Coefficient
AT1
AT2
Club
Early airway progenitor
Early stalk
Early tip
GHRL+ NE precursor
GHRL+ neuroendocrine
Interm neuroendocrine
Late airway progenitor
Late basal
Late stalk
Late tip
MUC5AC+ ASCL1+ progenitor
Mid airway progenitor
Mid basal
Mid stalk
Mid tip
Proximal basal
Proximal secretory 1
Proximal secretory 2
Proximal secretory 3
Proximal secretory progenitors
Pulmonary NE precursor
Pulmonary neuroendocrine
SMG
SMG basal
Squamous
matrix_62
p-LPC
d-LPC

**Fig. S1. Characterization of proximal and distal lung organoid differentiation. A**, Differentiation protocol workflow for generating proximal lung organoids (p-LOs) from hPSCs. **B**, Bright-field images comparing organoid morphology under untreated versus dissociation conditions during lung progenitor cell (LPC) induction. Scale bars, 500 µm (far left and middle two panels), 200 µm (right panels). Representative of three biologically independent experiments. **C**, Phase-contrast images of day 8 p-LOs and d-LOs counterstained with Hoechst. Scale bars, 500 µm. **D**, Organoid yield quantification (number per Matrigel droplet). **E**, Organoid diameter measurements. **F**, Representative flow cytometry analysis of NKX2-1 expression in p-LOs versus d-LOs. n = 3 independent experiments. **G-H**, Percentage (**G**) and mean fluorescence intensity (MFI) (**H**) of $NKX2\text{-}1^+$ cells. **I**, qPCR analysis of lung-specific and endodermal lineage markers in p-LOs and d-LOs derived from hESC lines (H1, H9) and induced pluripotent stem cells (hiPSCs). **J**, Integrated UMAP projection of p-LOs and d-LOs colored by annotated major cell types (left) and experimental conditions (right). **K**, Dot plot displaying canonical lineage markers for epithelial, mesenchymal, and neural cell populations. **L**, Proportional distribution of major cell types in d-LOs versus p-LOs. **M**, Epithelial subcluster marker expression distinguishing d-LPC, p-LPC, neuroendocrine cells, cycling cells, and hindgut-like cells. **N-O**, Gene Ontology (GO) enrichment analysis of biological processes enriched in p-LPC (**N**) and d-LPC (**O**) clusters. **P**, Expression patterns of proximal and distal lineage markers from fetal lung [15] in p-LPC and d-LPC clusters. **Q–S**, Heatmaps displaying the overlap coefficients of gene expression signatures from p-LPC and d-LPC clusters compared with cell type annotations from published human fetal lung scRNA-seq datasets by Cao et al., ref. 15 (**Q**), Sountoulidis et al., ref. 24 (**R**), and He et al., ref. 23 (**S**). **D**,**E**,**G**,**H**,**I**, Data are presented as mean ± SEM (n=3 biological replicates). *P*-values calculated using two-tailed Student's *t*-test.

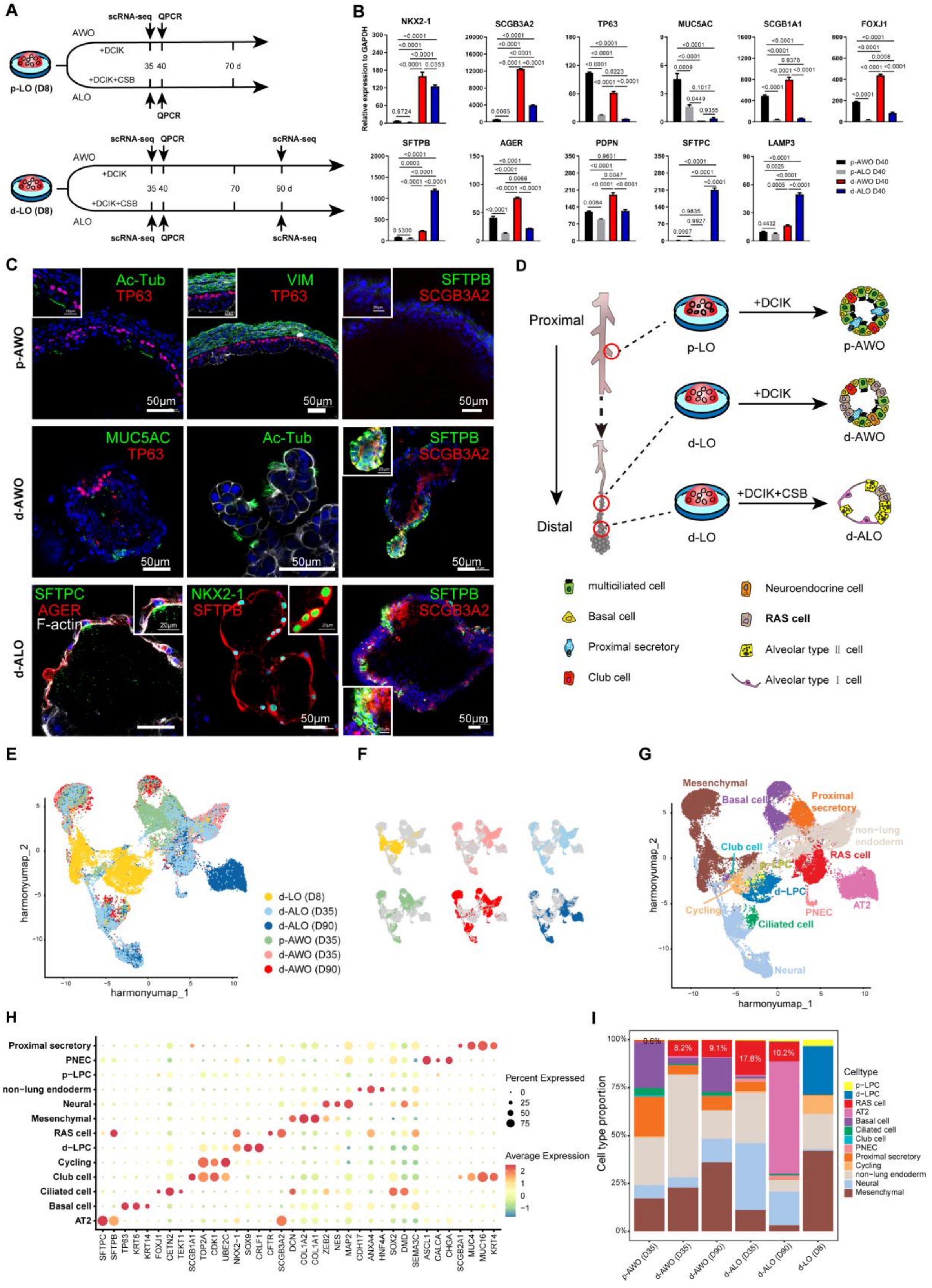
A
scRNA-seq
QPCR
AWO
+DCIK
+DCIK+CSB
ALO
35
40
70 d
p-LO (D8)
90 d
70
d-LO (D8)
B
NKX2-1
SCGB3A2
TP63
MUC5AC
SCGB1A1
FOXJ1
SFTPB
AGER
PDPN
SFTPC
LAMP3
Relative expression to GAPDH
p-AWO D40
p-ALO D40
d-AWO D40
d-ALO D40
C
p-AWO
d-AWO
d-ALO
Ac-Tub
TP63
VIM
SFTPB
SCGB3A2
MUC5AC
SFTPC
AGER
F-actin
NKX2-1
50μm
D
Proximal
Distal
p-LO
d-LO
+DCIK
+DCIK+CSB
p-AWO
d-AWO
d-ALO
multiciliated cell
Basal cell
Proximal secretory
Club cell
Neuroendocrine cell
RAS cell
Alveolar type Ⅱ cell
Alveolar type Ⅰ cell
E
harmonyumap_1
harmonyumap_2
d-LO (D8)
d-ALO (D35)
d-ALO (D90)
p-AWO (D35)
d-AWO (D35)
d-AWO (D90)
F
G
Mesenchymal
Basal cell
Proximal secretory
non-lung endoderm
Club cell
p-LPC
RAS cell
d-LPC
Cycling
PNEC
AT2
Ciliated cell
Neural
H
Proximal secretory
PNEC
p-LPC
non-lung endoderm
Neural
Mesenchymal
RAS cell
d-LPC
Cycling
Club cell
Ciliated cell
Basal cell
AT2
SFTPC
SFTPB
TP63
KRT5
KRT14
FOXJ1
CETN2
TEKT1
SCGB1A1
TOP2A
CDK1
UBE2C
NKX2-1
SOX9
CRLF1
CFTR
SCGB3A2
DCN
COL1A2
COL1A1
ZEB2
NES
MAP2
CDH17
ANXA4
HNF4A
SOX2
DMD
SEMA3C
ASCL1
CALCA
CHGA
SCGB2A1
MUC4
MUC16
KRT4
Percent Expressed
Average Expression
I
Cell type proportion
8.2%
9.1%
17.8%
10.2%
p-AWO (D35)
d-AWO (D35)
d-AWO (D90)
d-ALO (D35)
d-ALO (D90)
d-LO (D8)
Celltype
p-LPC
d-LPC
RAS cell
AT2
Basal cell
Ciliated cell
Club cell
PNEC
Proximal secretory
Cycling
non-lung endoderm
Neural
Mesenchymal

**Fig. S2. d-LOs generate distal airway and alveolar lineages containing RAS cells. A**, Differentiation timeline for generating airway and alveolar organoids from proximal (p-LO) and distal (d-LO) lung organoids. Arrows indicate time points for qPCR and scRNA-seq analyses. **B**, qPCR quantification of lung lineage markers in p-LO- and d-LO-derived organoids. Lung epithelial marker: *NKX2-1*; Airway markers: *SCGB3A2*, *TP63*, *MUC5AC*, *SCGB1A1*, *FOXJ1*; alveolar markers: *SFTPB*, *AGER*, *PDPN*, *SFTPC*, *LAMP3*. Data are presented as mean ± SEM (n=3). *P*-values calculated by one-way ANOVA with Tukey's multiple comparison test. **C**, Immunofluorescence co-staining of lineage markers in organoids: basal (TP63), distal secretory (SCGB3A2, SFTPB), multiciliated (Ac-Tub), mesenchymal (VIM), goblet (MUC5AC), AT2 (SFTPB, SFTPC, NKX2-1), and AT1 (AGER). Insets on select images highlight protein expression in specific cells. Scale bars, 50 µm (low-magnification images), 20 µm (expanded insets). Representative of three independent experiments. **D**, Schematic of regional lineage specification from proximal airway and distal airway/alveolar organoids. **E-F**, UMAP projections of single-cell RNA sequencing data from d-LOs and organoids under differentiation conditions, colored by all (**E**) or individual differentiation time/media conditions (**F**). **G**, Annotated UMAP distinguishing major cell types across differentiation conditions. **H**, Dot plot of canonical markers defining cell clusters in (**G**). **I**, Stacked bar plot quantifying the relative proportions of annotated cell types across different differentiation conditions.

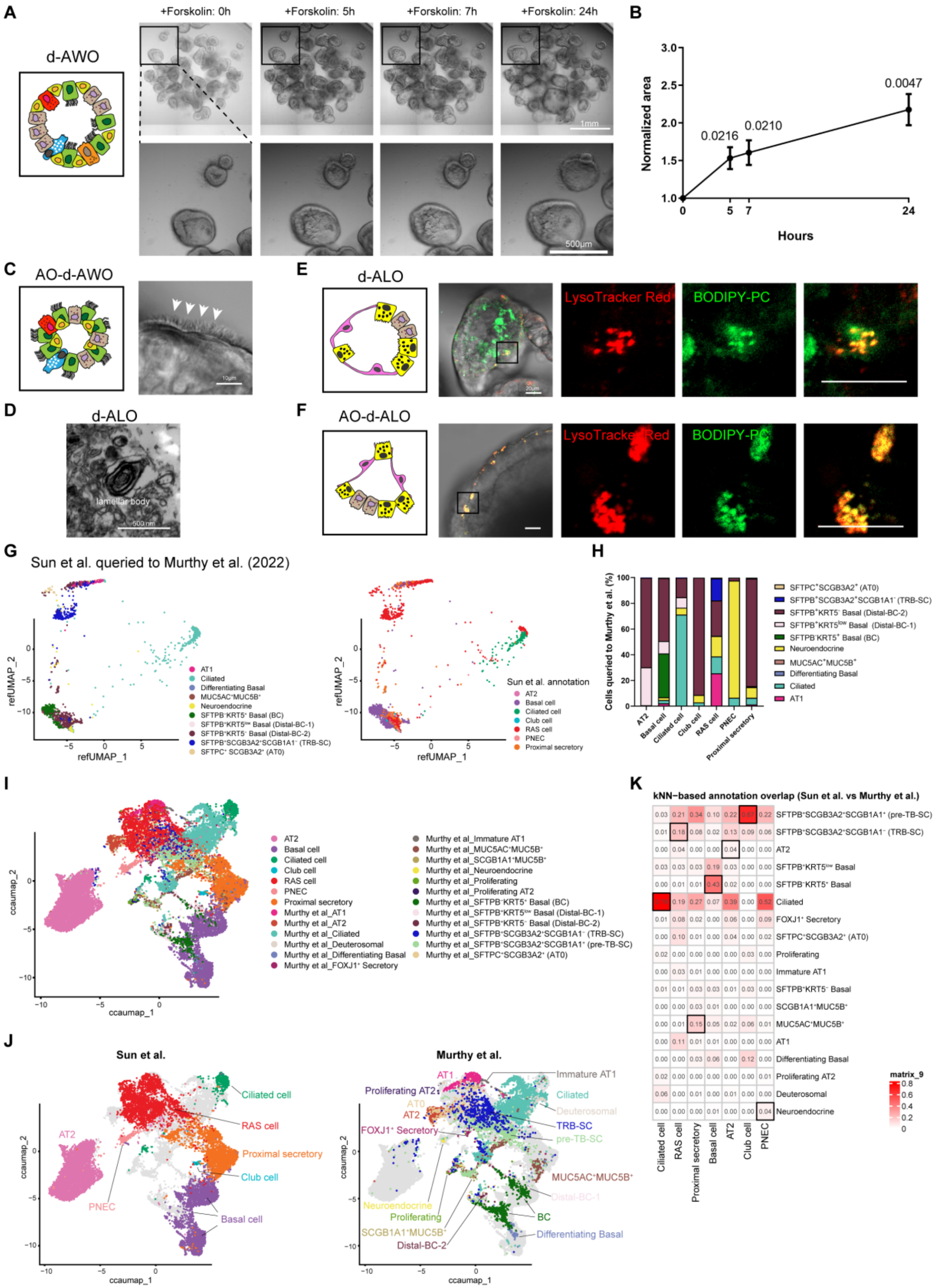
A
d-AWO
+Forskolin: 0h
+Forskolin: 5h
+Forskolin: 7h
+Forskolin: 24h
1mm
500μm
B
Normalized area
Hours
0.0216
0.0210
0.0047
C
AO-d-AWO
10μm
D
d-ALO
lamellar body
500 nm
E
d-ALO
LysoTracker Red
BODIPY-PC
F
AO-d-ALO
LysoTracker Red
BODIPY-PC
G
Sun et al. queried to Murthy et al. (2022)
refUMAP_1
refUMAP_2
AT1
Ciliated
Differentiating Basal
MUC5AC+MUC5B+
Neuroendocrine
SFTPB-KRT5+ Basal (BC)
SFTPB+KRT5low Basal (Distal-BC-1)
SFTPB+KRT5- Basal (Distal-BC-2)
SFTPB+SCGB3A2+SCGB1A1- (TRB-SC)
SFTPC+ SCGB3A2+ (AT0)
Sun et al. annotation
AT2
Basal cell
Ciliated cell
Club cell
RAS cell
PNEC
Proximal secretory
H
Cells queried to Murthy et al. (%)
I
ccaumap_1
ccaumap_2
Murthy et al_Immature AT1
Murthy et al_MUC5AC+MUC5B+
Murthy et al_SCGB1A1+MUC5B+
Murthy et al_Neuroendocrine
Murthy et al_Proliferating
Murthy et al_Proliferating AT2
Murthy et al_SFTPB-KRT5+ Basal (BC)
Murthy et al_SFTPB+KRT5low Basal (Distal-BC-1)
Murthy et al_SFTPB+KRT5- Basal (Distal-BC-2)
Murthy et al_SFTPB+SCGB3A2+SCGB1A1- (TRB-SC)
Murthy et al_SFTPB+SCGB3A2+SCGB1A1+ (pre-TB-SC)
Murthy et al_SFTPC+SCGB3A2+ (AT0)
Murthy et al_AT1
Murthy et al_AT2
Murthy et al_Ciliated
Murthy et al_Deuterosomal
Murthy et al_Differentiating Basal
Murthy et al_FOXJ1+ Secretory
J
Sun et al.
Murthy et al.
K
kNN−based annotation overlap (Sun et al. vs Murthy et al.)
SFTPB+SCGB3A2+SCGB1A1+ (pre-TB-SC)
SFTPB+SCGB3A2+SCGB1A1- (TRB-SC)
AT2
SFTPB+KRT5low Basal
SFTPB-KRT5+ Basal
Ciliated
FOXJ1+ Secretory
SFTPC+SCGB3A2+ (AT0)
Proliferating
Immature AT1
SFTPB+KRT5- Basal
SCGB1A1+MUC5B+
MUC5AC+MUC5B+
AT1
Differentiating Basal
Proliferating AT2
Deuterosomal
Neuroendocrine
matrix_9
Ciliated cell
RAS cell
Proximal secretory
Basal cell
AT2
Club cell
PNEC

**Fig. S3. Functional validation of distal lung organoid derivatives and cross-dataset comparison. A**, Time-lapse phase-contrast imaging of forskolin-induced swelling in distal airway organoids (d-AWO) derived from d-LOs. Scale bars, 1 mm (upper panel), 500 µm (lower panel). **B**, Quantification of normalized luminal swelling area in d-AWOs over 24 hours. Bars represent mean±SEM (n=20 organoids). *P*-values calculated using two-tailed Student's *t*-test. **C**, Schematic of apical-out (AO) polarity model (left) and representative microscopy showing cilia in AO-d-AWO. Scale bar, 10 µm. **D**, Transmission electron microscopy (TEM) of d-ALOs showing lamellar body-like structures. Scale bar, 500 nm. **E-F**, BODIPY-labeled phosphatidylcholine uptake in Matrigel-embedded (**E**) and apical-out (**F**) d-ALOs. Scale bars, 20 µm. Representative of three independent experiments. **G**, Reference mapping of Sun et al. scRNA-seq data onto the Murthy et al. (ref. 14) human fetal lung atlas using a query-to-reference projection. Left panel: cells colored by predicted cell types from Murthy et al.; Right panel: cells colored by Sun et al. original annotations. **H**, Stacked bar plot quantifying the composition of predicted Murthy et al. cell types within each annotated cluster from Sun et al. **I–J**, Integrated UMAP projection of combined datasets (**I**) and split views (**J**) showing the alignment of Sun et al. and Murthy et al. cells in a shared embedding space. **K**, Heatmap displaying kNN-based annotation overlap coefficients between Sun et al. clusters (columns) and Murthy et al. cell types (rows).

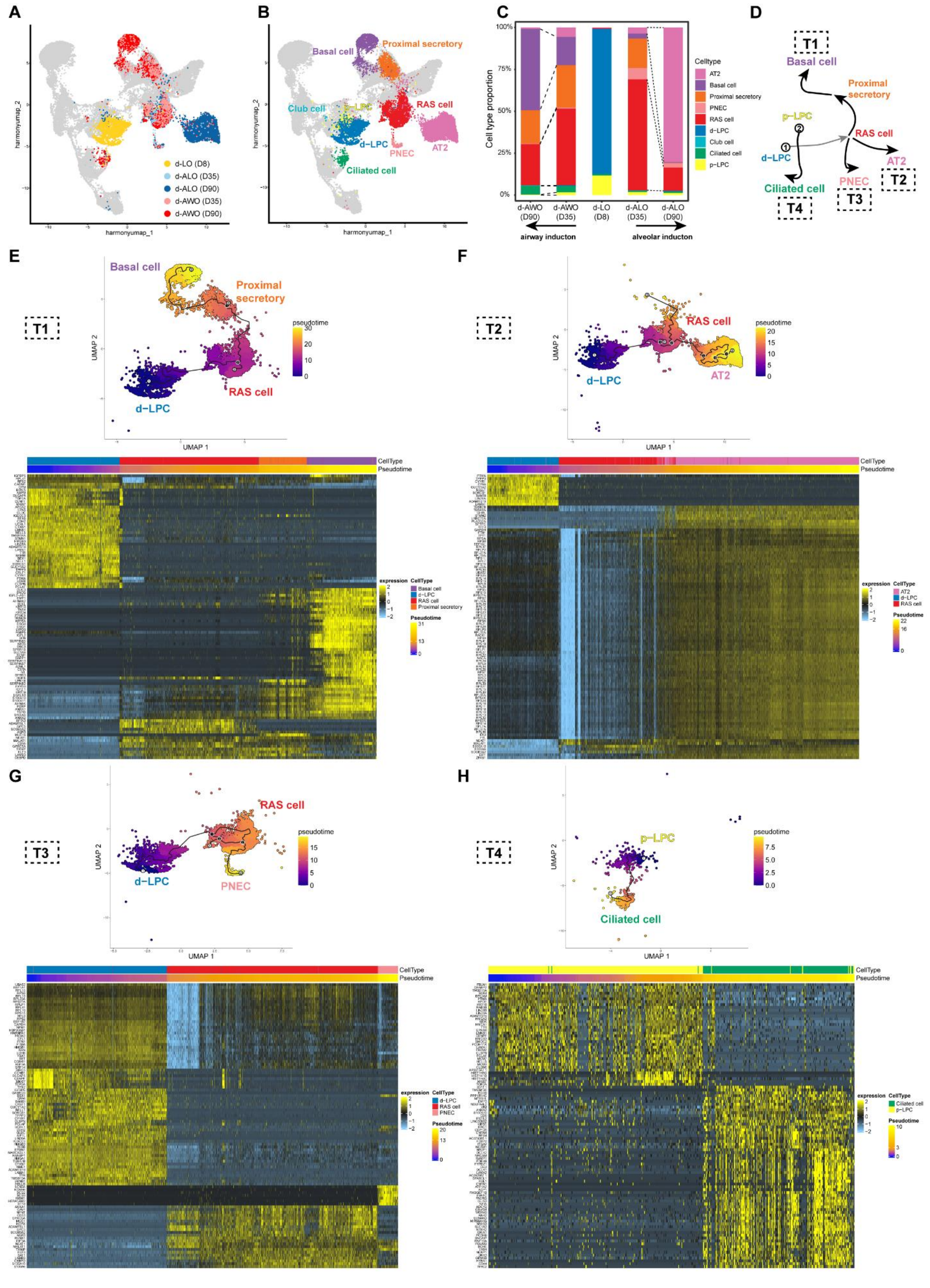
A
B
C
D
E
F
G
H
d-LO (D8)
d-ALO (D35)
d-ALO (D90)
d-AWO (D35)
d-AWO (D90)
harmonyumap_1
harmonyumap_2
Basal cell
Proximal secretory
Club cell
p-LPC
RAS cell
d-LPC
PNEC
AT2
Ciliated cell
Cell type proportion
Celltype
airway inducton
alveolar inducton
T1
T2
T3
T4
pseudotime
UMAP 1
UMAP 2
CellType
Pseudotime
expression

73

**Fig. S4. Lineage potential of RAS cells in d-LO-derived organoids. A-B**, UMAP projections of single-cell transcriptomes from d-LOs and their derivatives. Cells colored by experimental conditions (**A**) or annotated cell types (**B**). **C**, Proportional distribution of cell types in d-LOs versus d-LO-derived organoids. **D**, Schematic showing trajectory analysis identifying four differentiation paths from RAS cells to PNEC, basal, and AT2 lineages as well as from p-LPC to ciliated cell. **E-H**, UMAP projections depicting four cellular trajectories with color indicating progression (top) and heatmaps of trajectory-defining gene expression (bottom) across pseudotime for each lineage relationship.

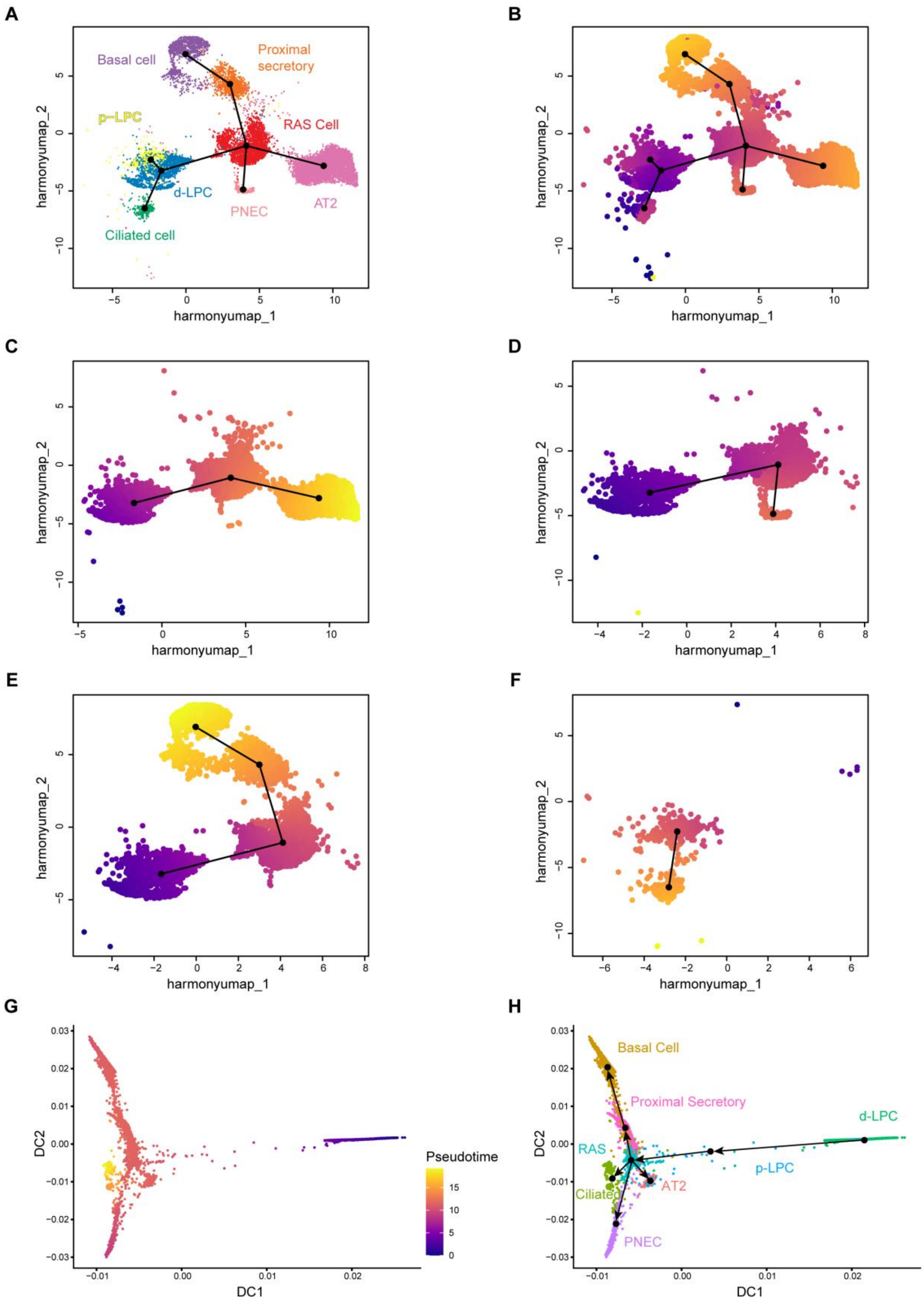
A
B
C
D
E
F
G
H
Basal cell
Proximal secretory
p-LPC
RAS Cell
d-LPC
PNEC
AT2
Ciliated cell
harmonyumap_1
harmonyumap_2
DC1
DC2
Pseudotime
Basal Cell
Proximal Secretory
d-LPC
RAS
p-LPC
Ciliated
AT2
PNEC

**Fig. S5. Independent validation of lineage trajectory using Slingshot and diffusion pseudotime analyses.** **A**, UMAP projection of integrated d-LO and d-LO-derived organoid epithelial cells with Slingshot-inferred lineage trajectories overlaid. Cell types are annotated by color. **B**, Pseudotemporal ordering of epithelial subtypes along the Slingshot-derived trajectory. Cells are colored by pseudotime progression. **C**-**F**, UMAP projections depicting four distinct differentiation trajectories with color indicating progression for each lineage relationship. **G**, Diffusion pseudotime (DPT) analysis of integrated epithelial populations. Cells colored according to diffusion pseudotime values, showing the global trajectory progression. **H**, The same DPT embedding colored by annotated cell types. Arrows indicate the predicted developmental progression.

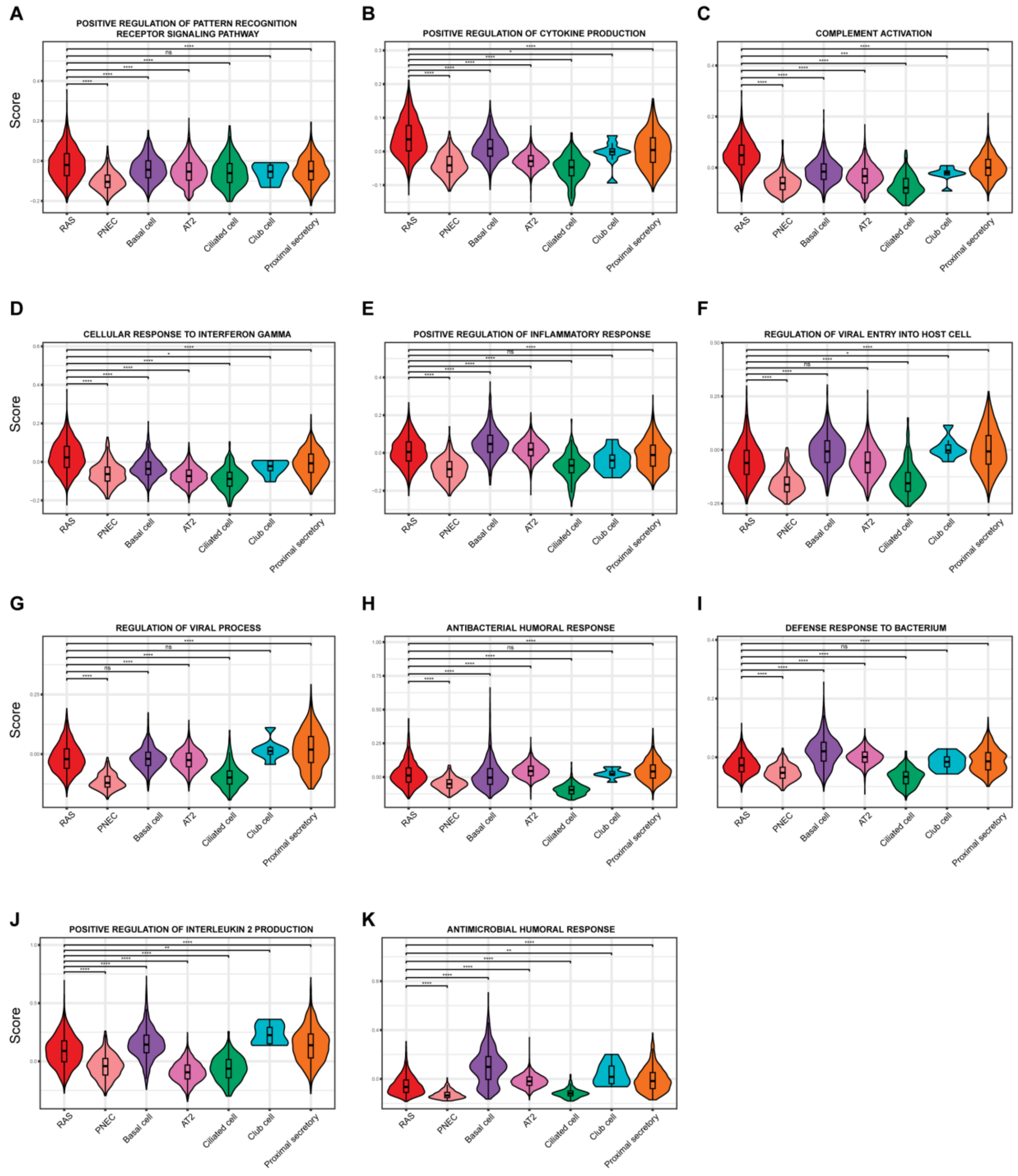
A
POSITIVE REGULATION OF PATTERN RECOGNITION RECEPTOR SIGNALING PATHWAY
B
POSITIVE REGULATION OF CYTOKINE PRODUCTION
C
COMPLEMENT ACTIVATION
D
CELLULAR RESPONSE TO INTERFERON GAMMA
E
POSITIVE REGULATION OF INFLAMMATORY RESPONSE
F
REGULATION OF VIRAL ENTRY INTO HOST CELL
G
REGULATION OF VIRAL PROCESS
H
ANTIBACTERIAL HUMORAL RESPONSE
I
DEFENSE RESPONSE TO BACTERIUM
J
POSITIVE REGULATION OF INTERLEUKIN 2 PRODUCTION
K
ANTIMICROBIAL HUMORAL RESPONSE
Score
RAS
PNEC
Basal cell
AT2
Ciliated cell
Club cell
Proximal secretory

**Fig. S6. Enrichment of immune- and host defense–associated pathways across epithelial populations in organoids. A-K**, Violin plots showing module scores for selected immune- and host defense–related pathways across major epithelial cell populations. Pathways analyzed include positive regulation of pattern-recognition receptor signaling (**A**), positive regulation of cytokine production (**B**), complement activation (**C**), cellular response to interferon-γ (**D**), positive regulation of inflammatory response (**E**), regulation of viral entry into host cells (**F**), regulation of viral processes (**G**), antibacterial humoral response (**H**), defense response to bacterium (**I**), positive regulation of interleukin-2 production (**J**), and antimicrobial humoral response (**K**). RAS cells showed prominent enrichment of multiple immune-sensing, inflammatory, and host defense programs.

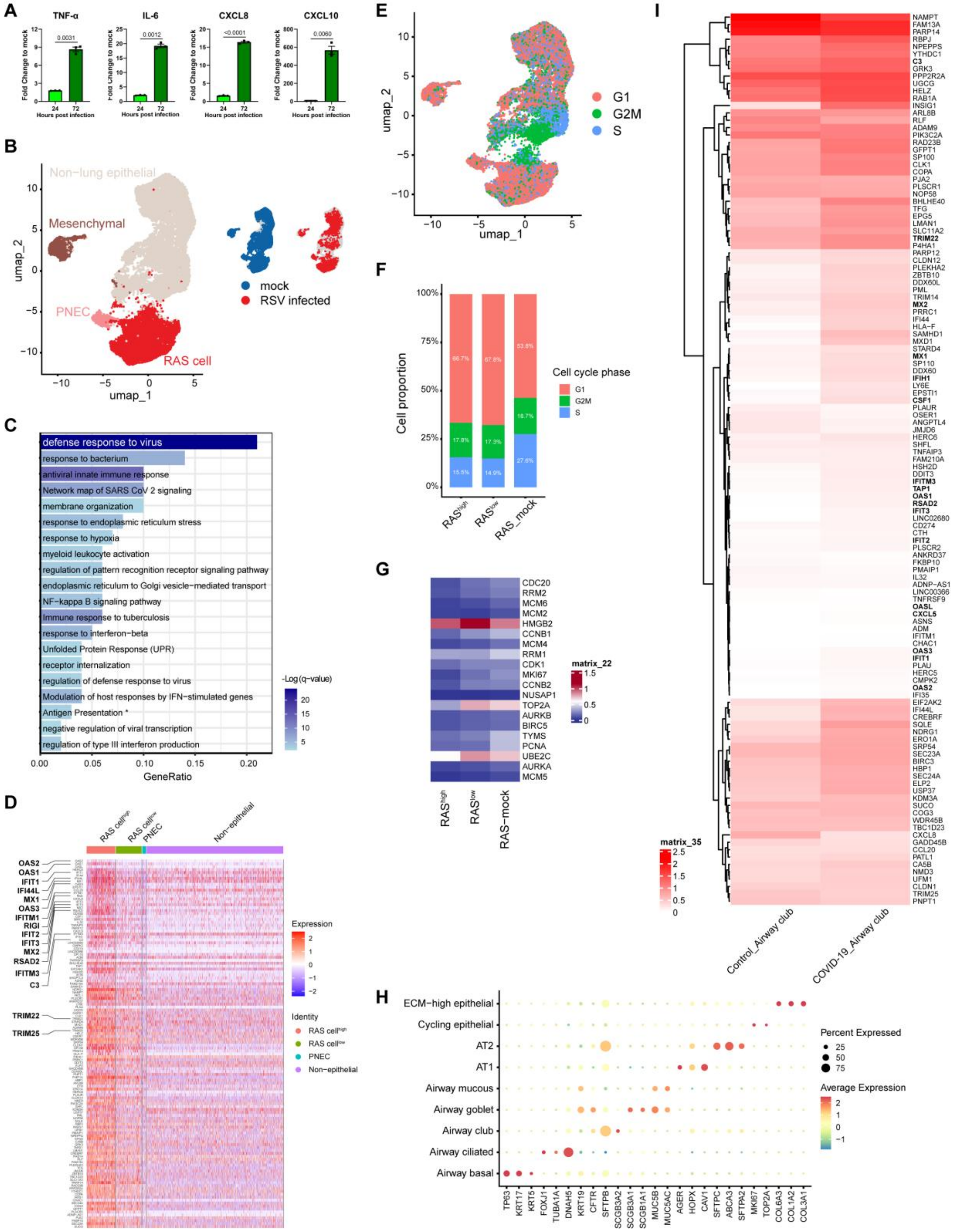
A
TNF-α
IL-6
CXCL8
CXCL10
Fold Change to mock
Hours post infection
0.0031
0.0012
<0.0001
0.0060
B
Non−lung epithelial
Mesenchymal
PNEC
RAS cell
mock
RSV infected
umap_1
umap_2
C
defense response to virus
response to bacterium
antiviral innate immune response
Network map of SARS CoV 2 signaling
membrane organization
response to endoplasmic reticulum stress
response to hypoxia
myeloid leukocyte activation
regulation of pattern recognition receptor signaling pathway
endoplasmic reticulum to Golgi vesicle−mediated transport
NF−kappa B signaling pathway
Immune response to tuberculosis
response to interferon−beta
Unfolded Protein Response (UPR)
receptor internalization
regulation of defense response to virus
Modulation of host responses by IFN−stimulated genes
Antigen Presentation *
negative regulation of viral transcription
regulation of type III interferon production
-Log(q-value)
GeneRatio
D
RAS cell^high
RAS cell^low
PNEC
Non−epithelial
OAS2
OAS1
IFIT1
IFI44L
MX1
OAS3
IFITM1
RIGI
IFIT2
IFIT3
MX2
RSAD2
IFITM3
C3
TRIM22
TRIM25
Expression
Identity
E
G1
G2M
S
F
Cell proportion
Cell cycle phase
RAS^high
RAS^low
RAS_mock
G
CDC20
RRM2
MCM6
MCM2
HMGB2
CCNB1
MCM4
RRM1
CDK1
MKI67
CCNB2
NUSAP1
TOP2A
AURKB
BIRC5
TYMS
PCNA
UBE2C
AURKA
MCM5
matrix_22
RAS-mock
H
ECM−high epithelial
Cycling epithelial
AT2
AT1
Airway mucous
Airway goblet
Airway club
Airway ciliated
Airway basal
Percent Expressed
Average Expression
I
matrix_35
Control_Airway club
COVID-19_Airway club

**Fig. S7. Immune defense capacity of RAS cells in d-LO derived organoids infected by RSV.** **A**, qPCR analysis of *CXCL8*, *CXCL10*, *IL-6*, and *TNF-α* expression in d-ALOs infected with RSV (MOI 0.1) at 24 and72 hours post-infection (hpi). Data are presented as mean ± SEM (n=3). *P*-values calculated using two-tailed Student's *t*-test. **B**, Integrated UMAP projection of scRNA-seq data from mock- and RSV-infected organoids, colored by annotated cell types (left) and infection status (right). **C**, Gene Ontology (GO) enrichment analysis of shared DE genes, highlighting "defense response to virus" as the top pathway. The full name of the term marked by an asterisk is 'Antigen Presentation: Folding, assembly and peptide loading of class I MHC'. **D**, Heatmap of 126 common DE genes across clusters in RSV-infected d-ALOs. **E**–**G**, Cell cycle analysis of scRNA-seq data from mock- and RSV-infected organoids. UMAP projection colored by cell cycle phase (**E**), stacked bar plot quantifying the proportion of cells in G1, G2M, and S phases (**F**), and heatmap of cell cycle regulatory genes (**G**) in $RAS^{high}$ cell, $RAS^{low}$ cell, and RAS-mock groups. **H**, Dot plot showing the expression of canonical lineage markers identifying distinct cell populations in the single-cell lung atlas of lethal COVID-19 (Melms et al., ref. 32). **I**, Heatmap validating the expression of the 121 core antiviral signature genes in airway club cells from control and COVID-19 patients (Melms et al., ref. 32).

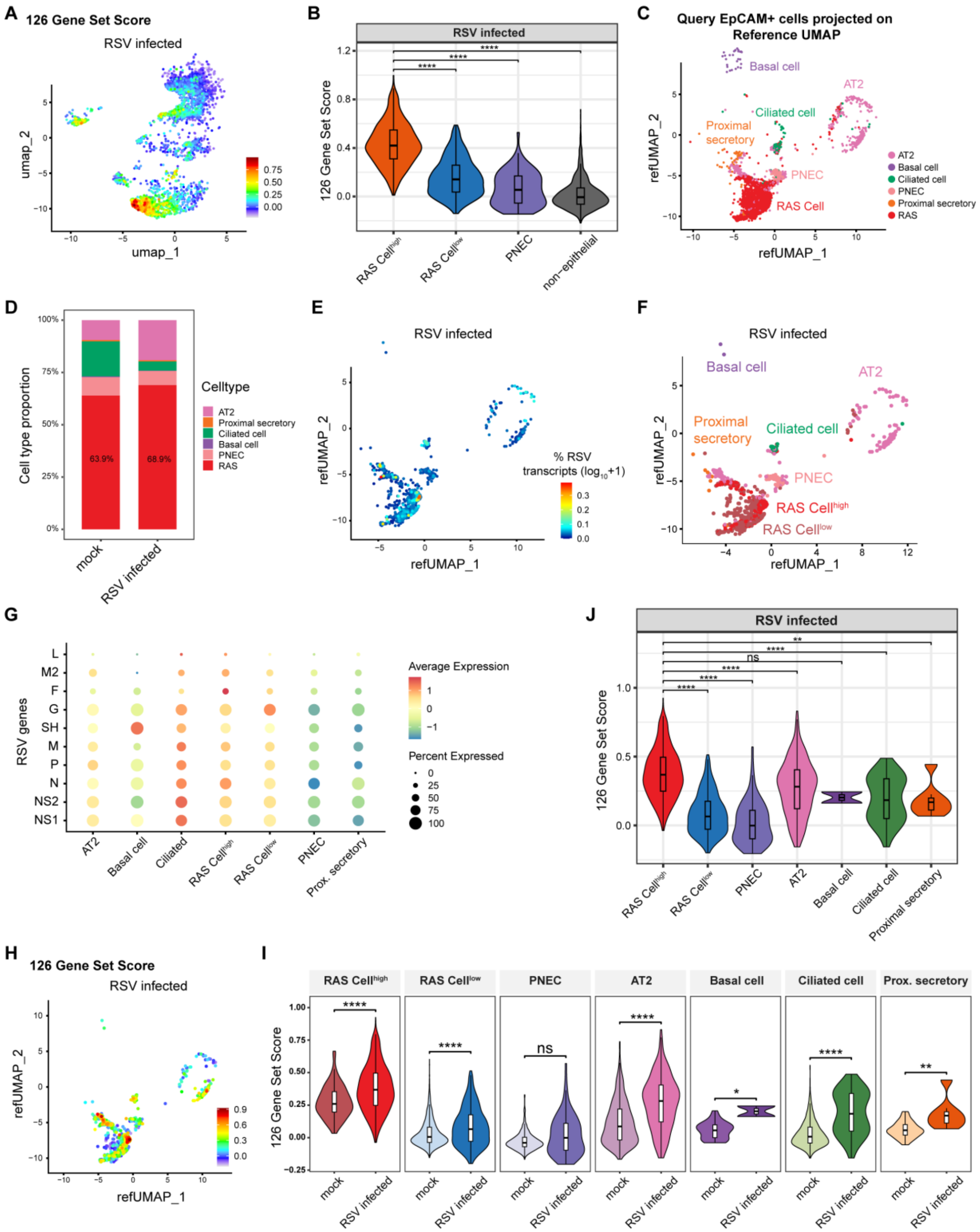
A
126 Gene Set Score
RSV infected
umap_1
umap_2
B
RSV infected
126 Gene Set Score
RAS Cellhigh
RAS Celllow
PNEC
non-epithelial
C
Query EpCAM+ cells projected on Reference UMAP
refUMAP_1
refUMAP_2
Basal cell
AT2
Ciliated cell
Proximal secretory
PNEC
RAS Cell
RAS
D
Cell type proportion
Celltype
AT2
Proximal secretory
Ciliated cell
Basal cell
PNEC
RAS
63.9%
68.9%
mock
RSV infected
E
RSV infected
% RSV transcripts (log10+1)
F
RSV infected
RAS Cellhigh
RAS Celllow
G
RSV genes
L
M2
F
G
SH
M
P
N
NS2
NS1
Average Expression
Percent Expressed
AT2
Basal cell
Ciliated
Prox. secretory
J
RSV infected
ns
Proximal secretory
H
126 Gene Set Score
RSV infected
I
Prox. secretory
mock
RSV infected

**Fig. S8. Cell-type–resolved analysis of RSV infection responses across epithelial populations.** **A**, UMAP projection of RSV-infected cells colored by the 126 RSV-response gene signature score. **B**, Violin plot comparing RSV-response signature scores among RAS cells, PNECs, and non-epithelial populations. **C**, Reference-based mapping of $EpCAM^{+}$ cells onto the integrated distal lung epithelial atlas. **D**, Cell-type composition of mock and RSV-infected organoid cultures. **E**–**G**, Distribution of RSV transcripts across epithelial populations. UMAP projection colored by RSV transcript abundance (**E**), identification of RAS $Cell^{high}$ and RAS $Cell^{low}$ subsets (**F**), and dot plot showing expression of RSV genes across epithelial cell types (**G**). **H**. UMAP projection colored by the 126 RSV-response signature score. **I**. Comparison of signature scores between mock and RSV-infected conditions across epithelial populations. **J**. Comparison of RSV-response signature scores among epithelial cell types in infected cultures.

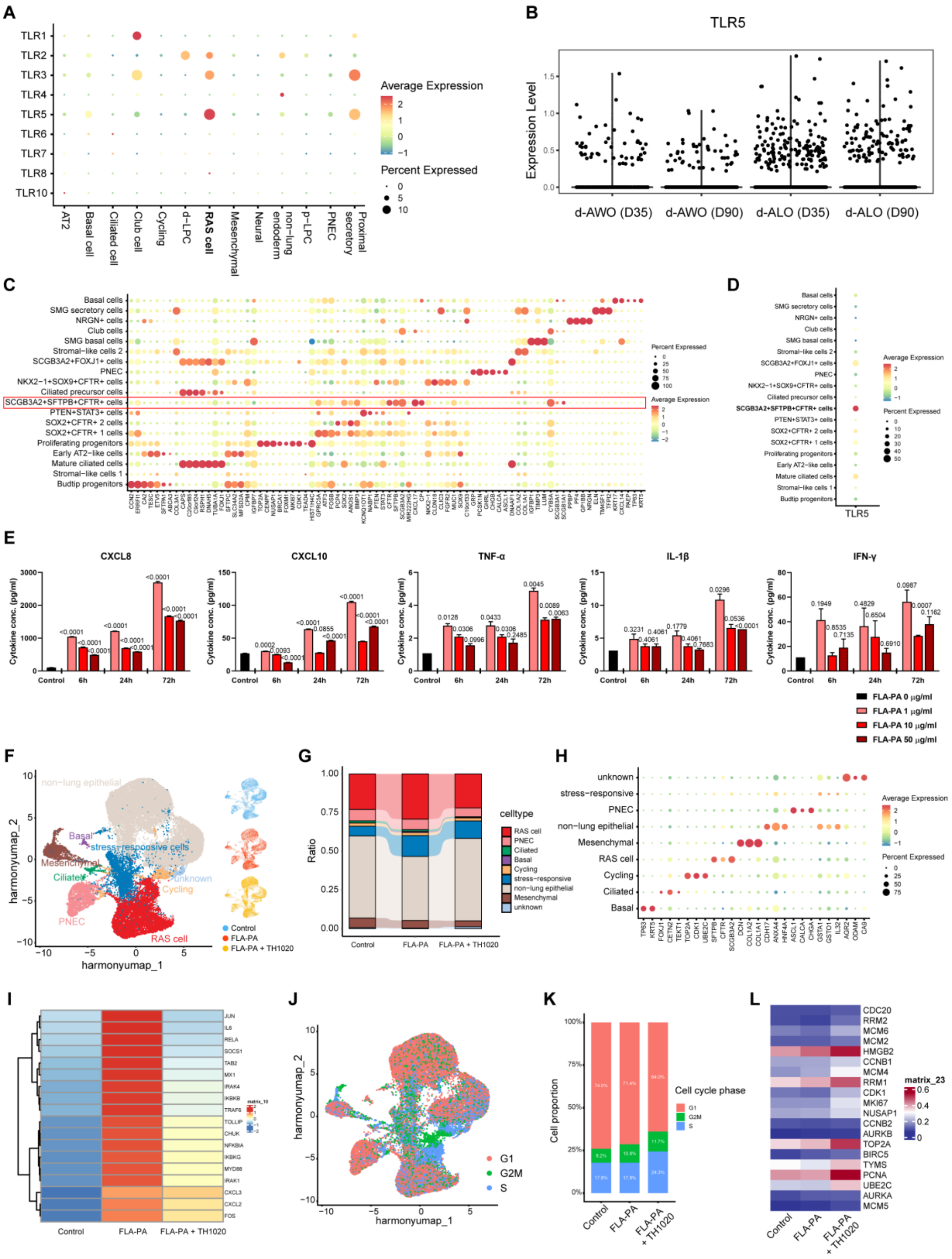
A
TLR1
TLR2
TLR3
TLR4
TLR5
TLR6
TLR7
TLR8
TLR10
Average Expression
Percent Expressed
AT2
Basal cell
Ciliated cell
Club cell
Cycling
d-LPC
RAS cell
Mesenchymal
Neural
non-lung endoderm
p-LPC
PNEC
Proximal secretory
B
TLR5
Expression Level
d-AWO (D35)
d-AWO (D90)
d-ALO (D35)
d-ALO (D90)
C
Basal cells
SMG secretory cells
NRGN+ cells
Club cells
SMG basal cells
Stromal-like cells 2
SCGB3A2+FOXJ1+ cells
PNEC
NKX2-1+SOX9+CFTR+ cells
Ciliated precursor cells
SCGB3A2+SFTPB+CFTR+ cells
PTEN+STAT3+ cells
SOX2+CFTR+ 2 cells
SOX2+CFTR+ 1 cells
Proliferating progenitors
Early AT2-like cells
Mature ciliated cells
Stromal-like cells 1
Budtip progenitors
Percent Expressed
Average Expression
D
TLR5
E
CXCL8
CXCL10
TNF-α
IL-1β
IFN-γ
Cytokine conc. (pg/ml)
Control
6h
24h
72h
FLA-PA 0 μg/ml
FLA-PA 1 μg/ml
FLA-PA 10 μg/ml
FLA-PA 50 μg/ml
F
harmonyumap_2
harmonyumap_1
non-lung epithelial
Basal
stress-responsive cells
Mesenchymal
Ciliated
unknown
Cycling
PNEC
RAS cell
Control
FLA-PA
FLA-PA + TH1020
G
Ratio
celltype
RAS cell
PNEC
Ciliated
Basal
Cycling
stress-responsive
non-lung epithelial
Mesenchymal
unknown
Control
FLA-PA
FLA-PA + TH1020
H
unknown
stress-responsive
PNEC
non-lung epithelial
Mesenchymal
RAS cell
Cycling
Ciliated
Basal
Average Expression
Percent Expressed
I
Control
FLA-PA
FLA-PA + TH1020
J
harmonyumap_2
harmonyumap_1
G1
G2M
S
K
Cell proportion
Cell cycle phase
G1
G2M
S
Control
FLA-PA
FLA-PA + TH1020
L
CDC20
RRM2
MCM6
MCM2
HMGB2
CCNB1
MCM4
RRM1
CDK1
MKI67
NUSAP1
CCNB2
AURKB
TOP2A
BIRC5
TYMS
PCNA
UBE2C
AURKA
MCM5
matrix_23
Control
FLA-PA
FLA-PA + TH1020

**Fig. S9. scRNA-seq analysis of RAS cells upon TLR5 activation.** **A**, Dot plot showing the expression of Toll-like receptors (TLRs) across all cell clusters (Fig. S2H). **B**, Violin plots comparing TLR5 expression in RAS cells under indicated experimental conditions. **C**–**D**, Validation of cell type markers and TLR5 expression in the human fetal lung atlas from Quach et al. (ref. 34). **C**, Dot plot of lineage markers identifying distinct cell populations; the red box highlights the gene signature of $SCGB3A2^{+}SFTPB^{+}CFTR^{+}$ cells. **D**, Dot plot displaying TLR5 expression levels across the cell types defined in (**C**). **E**, Cytokine/chemokine secretion (CXCL8, CXCL10, IL-6, IL-1β, TNF-α) from d-ALOs stimulated with increasing concentrations of FLA-PA protein (1–50 µg/ml) over 6–72 hours. Data are presented as mean ± SEM (n=3 biological replicates). *P*-values calculated using two-way ANOVA. **F**, UMAP projections of single-cell transcriptomes from control, FLA-PA-treated, and FLA-PA+TH1020-treated d-ALOs. Cells colored by experimental conditions (right) or annotated cell types (left). **G**, Proportional distribution of cell types in control, FLA-PA-treated, and FLA-PA+TH1020-treated d-ALOs. **H**, Cluster defining genes within (**F**) are shown in dot plot format. **I**, Heatmap from pseudo-bulk analysis of scRNA-seq data showing changes in mRNA expression of TLR signaling pathway related genes in RAS cells stimulated with FLA-PA (1µg/ml) with or without TH1020 (1µM) for 72h. **J**–**L**, Cell cycle analysis of the cell populations shown in (**F**). UMAP colored by cell cycle phase (**J**), stacked bar plot quantifying phase proportions (**K**) and heatmap of cell cycle regulatory genes (**L**) in RAS cell cluster across treatment groups.

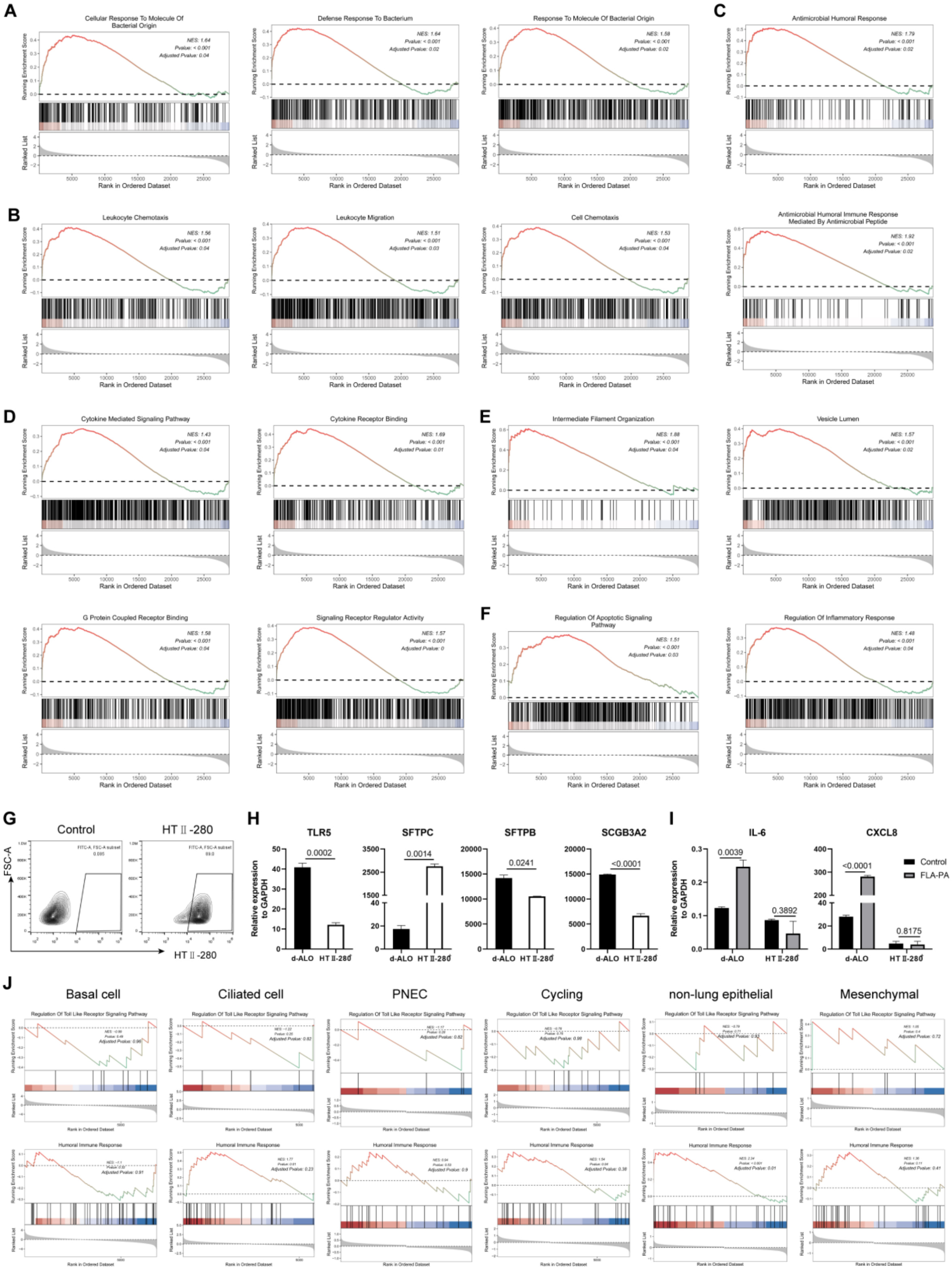
A
Cellular Response To Molecule Of Bacterial Origin
Defense Response To Bacterium
Response To Molecule Of Bacterial Origin
C
Antimicrobial Humoral Response
B
Leukocyte Chemotaxis
Leukocyte Migration
Cell Chemotaxis
Antimicrobial Humoral Immune Response Mediated By Antimicrobial Peptide
D
Cytokine Mediated Signaling Pathway
Cytokine Receptor Binding
E
Intermediate Filament Organization
Vesicle Lumen
G Protein Coupled Receptor Binding
Signaling Receptor Regulator Activity
F
Regulation Of Apoptotic Signaling Pathway
Regulation Of Inflammatory Response
Running Enrichment Score
Ranked List
Rank in Ordered Dataset
G
Control
HT II -280
FSC-A
HT II -280
H
TLR5
SFTPC
SFTPB
SCGB3A2
Relative expression to GAPDH
0.0002
0.0014
0.0241
<0.0001
d-ALO
HT II-280
I
IL-6
CXCL8
0.0039
0.3892
<0.0001
0.8175
Control
FLA-PA
J
Basal cell
Ciliated cell
PNEC
Cycling
non-lung epithelial
Mesenchymal
Regulation Of Toll Like Receptor Signaling Pathway
Humoral Immune Response

**Fig. S10. Specificity of FLA-PA-induced immune responses in RAS cells. A-F**, GSEA reveals significantly upregulated items in RAS cells after stimulated with FLA-PA, indicating that these cells activated a series of functions related to immune recognition (**A**), leukocyte recruitment (**B**), antibacterial response (**C**), cytokine and receptor signal regulation (**D**), cytoskeleton remodeling and secretion regulation (**E**), and programmed cell death and immune regulation (**F**). **G**, Flow cytometry gating strategy for AT2 cell isolation using HTII-280 surface marker. **H**, qPCR validation of *TLR5*, *SFTPC*, *SFTPB*, and *SCGB3A2* expression in sorted HTII-280+ AT2 cells, confirming high alveolar (SFTPC) and low immune (TLR5) signatures. **I**, Comparative RT-PCR analysis of *CXCL8* and *IL-6* in whole d-ALOs versus sorted HTII-280+ AT2-enriched organoids after 72 h FLA-PA stimulation (1 μg/ml). **J**, GSEA plots evaluating "Regulation of Toll Like Receptor Signaling Pathway" (top row) and "Humoral Immune Response" (bottom row) signatures across other cell types (Basal, Ciliated, PNEC, Cycling, non-lung epithelial, Mesenchymal). **H**,**I**, Data are presented as mean ± SEM (n=3). *P*-values calculated using two-tailed Student's *t*-test.

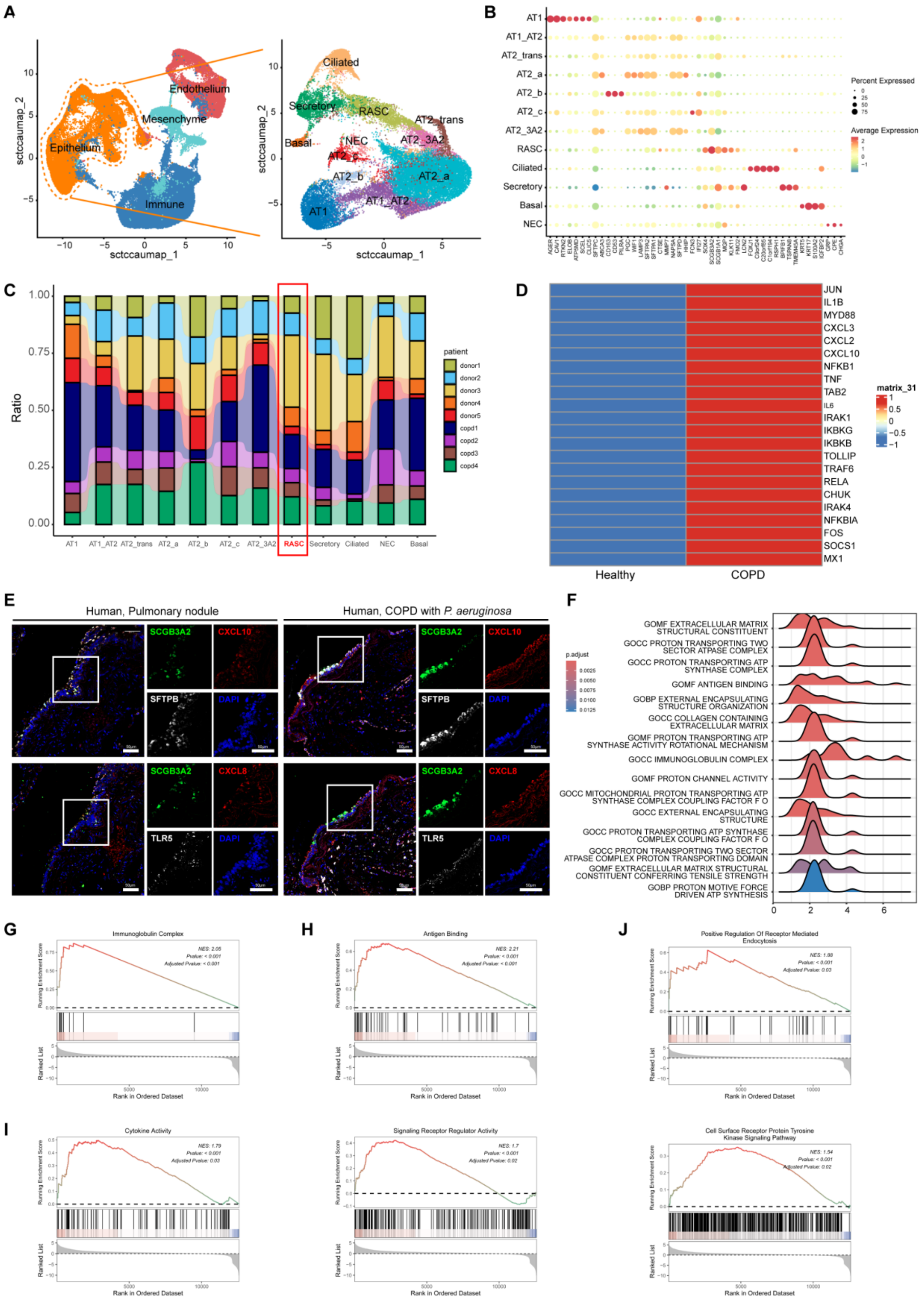
A
Endothelium
Mesenchyme
Epithelium
Immune
sctccaumap_2
sctccaumap_1
Ciliated
Secretory
RASC
AT2_trans
Basal
NEC
AT2_3A2
AT2_c
AT2_b
AT2_a
AT1
AT1_AT2
B
Percent Expressed
Average Expression
C
Ratio
patient
donor1
donor2
donor3
donor4
donor5
copd1
copd2
copd3
copd4
D
JUN
IL1B
MYD88
CXCL3
CXCL2
CXCL10
NFKB1
TNF
TAB2
IL6
IRAK1
IKBKG
IKBKB
TOLLIP
TRAF6
RELA
CHUK
IRAK4
NFKBIA
FOS
SOCS1
MX1
matrix_31
Healthy
COPD
E
Human, Pulmonary nodule
Human, COPD with P. aeruginosa
SCGB3A2
CXCL10
SFTPB
DAPI
CXCL8
TLR5
F
p.adjust
GOMF EXTRACELLULAR MATRIX STRUCTURAL CONSTITUENT
GOCC PROTON TRANSPORTING TWO SECTOR ATPASE COMPLEX
GOCC PROTON TRANSPORTING ATP SYNTHASE COMPLEX
GOMF ANTIGEN BINDING
GOBP EXTERNAL ENCAPSULATING STRUCTURE ORGANIZATION
GOCC COLLAGEN CONTAINING EXTRACELLULAR MATRIX
GOMF PROTON TRANSPORTING ATP SYNTHASE ACTIVITY ROTATIONAL MECHANISM
GOCC IMMUNOGLOBULIN COMPLEX
GOMF PROTON CHANNEL ACTIVITY
GOCC MITOCHONDRIAL PROTON TRANSPORTING ATP SYNTHASE COMPLEX COUPLING FACTOR F O
GOCC EXTERNAL ENCAPSULATING STRUCTURE
GOCC PROTON TRANSPORTING ATP SYNTHASE COMPLEX COUPLING FACTOR F O
GOCC PROTON TRANSPORTING TWO SECTOR ATPASE COMPLEX PROTON TRANSPORTING DOMAIN
GOMF EXTRACELLULAR MATRIX STRUCTURAL CONSTITUENT CONFERRING TENSILE STRENGTH
GOBP PROTON MOTIVE FORCE DRIVEN ATP SYNTHESIS
G
Immunoglobulin Complex
H
Antigen Binding
J
Positive Regulation Of Receptor Mediated Endocytosis
I
Cytokine Activity
Signaling Receptor Regulator Activity
Cell Surface Receptor Protein Tyrosine Kinase Signaling Pathway
Running Enrichment Score
Ranked List
Rank in Ordered Dataset

**Fig. S11. scRNA-seq analysis of RAS cells from COPD. A**, Reanalysis of scRNAseq dataset from normal and COPD peripheral samples (Basil *et al.*, ref. 13) and subset of epithelium showing expected epithelial populations. The COPD cohort consists of patients with severe COPD as defined by FEV1 values from the source study (Basil et al., ref. 13), including one patient with GOLD stage III and three patients with GOLD stage IV COPD. **B**, Cluster defining genes within the epithelium are shown in dot plot format. **C**, Stacked bar graphs showing the normal and COPD patients contribution to each epithelial cell cluster. **D**, Heatmap from pseudo-bulk analysis of scRNA-seq data showing significant changes in mRNA expression of TLR signaling pathway related genes in RAS cells from normal and COPD donors. **E**, Immunofluorescence of SCGB3A2 (green), CXCL8/CXCL10 (red), SFTPB/TLR5 (white), and DAPI (blue) in human lung sections from control and *Pseudomonas aeruginosa*-infected COPD patients. White boxed area is expanded. Scale bars, 50 µm. **F**, Top 15 GO terms enriched in DEGs between RAS cells from COPD versus from healthy controls. Significantly enriched terms highlight increased mitochondrial ATP synthesis, extracellular matrix organization, and immune-related processes, suggesting functional adaptation of RAS cells under chronic disease conditions. **G-J**, GSEA reveals significantly upregulated immune related items in RAS cells from COPD, including antibody-related immune function (**G**), antigen recognition (**H**), Cytokine signaling and immune regulation (**I**), and endocytosis (**J**).

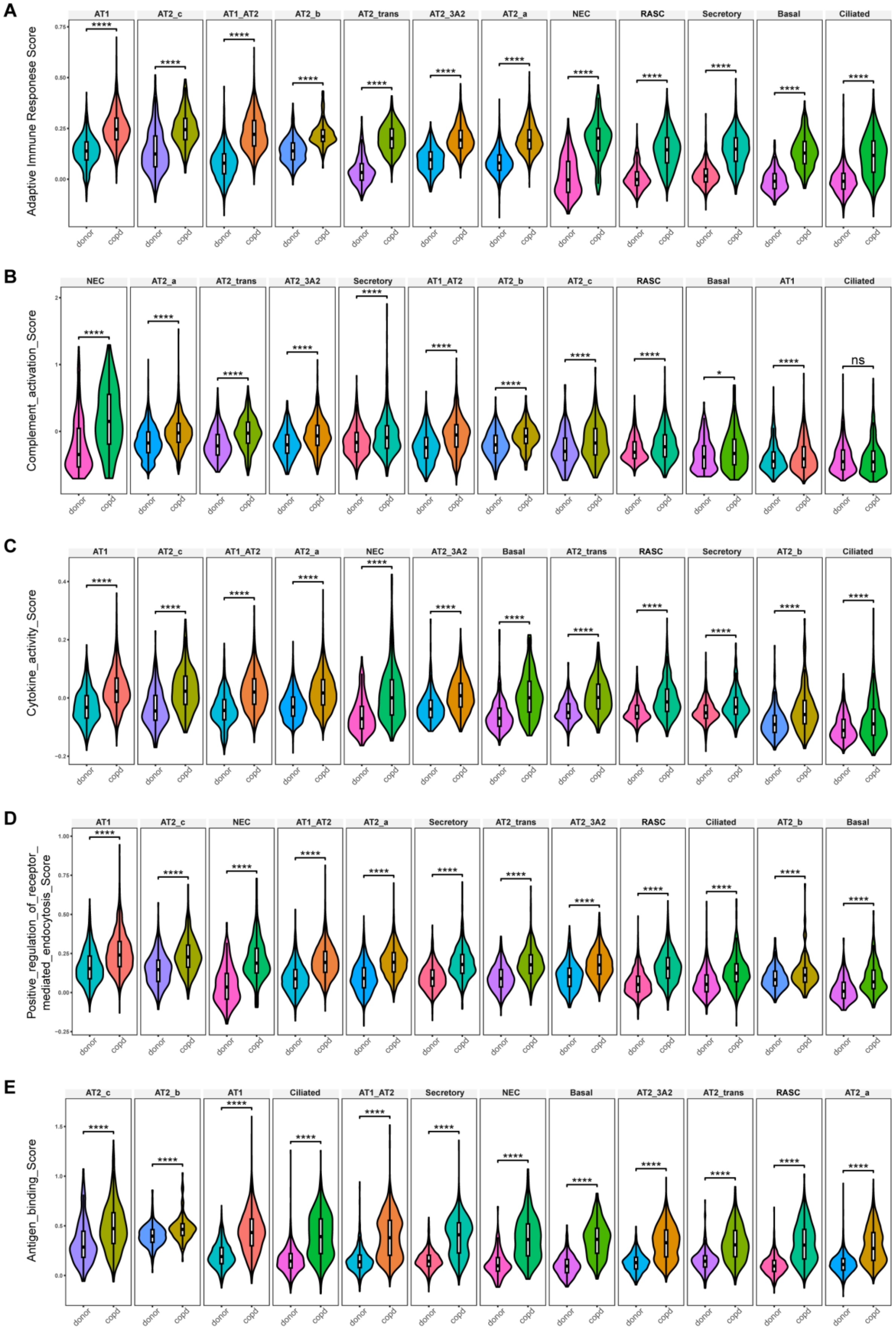
A
Adaptive Immune Responese Score
AT1
AT2_c
AT1_AT2
AT2_b
AT2_trans
AT2_3A2
AT2_a
NEC
RASC
Secretory
Basal
Ciliated
B
Complement_activation_Score
NEC
AT2_a
AT2_trans
AT2_3A2
Secretory
AT1_AT2
AT2_b
AT2_c
RASC
Basal
AT1
Ciliated
ns
C
Cytokine_activity_Score
AT1
AT2_c
AT1_AT2
AT2_a
NEC
AT2_3A2
Basal
AT2_trans
RASC
Secretory
AT2_b
Ciliated
D
Positive_regulation_of_receptor_
mediated_endocytosis_Score
AT1
AT2_c
NEC
AT1_AT2
AT2_a
Secretory
AT2_trans
AT2_3A2
RASC
Ciliated
AT2_b
Basal
E
Antigen_binding_Score
AT2_c
AT2_b
AT1
Ciliated
AT1_AT2
Secretory
NEC
Basal
AT2_3A2
AT2_trans
RASC
AT2_a
donor
copd

214

**Fig. S12. Comparison of immune-associated pathway activities across epithelial populations in healthy and COPD lungs. A-E**, Violin plots showing gene signature enrichment scores for representative immune-related pathways across major epithelial cell populations from healthy donor and COPD lungs. Pathways analyzed include adaptive immune response (**A**), complement activation (**B**), cytokine activity (**C**), positive regulation of receptor-mediated endocytosis (**D**), and antigen binding (**E**).

**Table S1.**

All qPCR primers used in the described experiments.

| qRT-PCR primers | | |
|---|---|---|
| GENE | Forward 5' - 3' | Reverse 5' - 3' |
| GAPDH | ACAACTTTGGTATCGTGGAAGG | GCCATCACGCCACAGTTTC |
| NKX2-1 | CTCATGTTCATGCCGCTC | GACACCATGAGGAACAGCG |
| SOX2 | GCTTAGCCTCGTCGATGAAC | AACCCCAAGATGCACAACTC |
| SOX9 | GTACCCGCACTTGCACAAC | GTGGtCCTTCTTGTGCTGC |
| PAX6 | CGAATTCTGCAGGTGTCCAA | ACAGACCCCCTCGGACAGTAAT |
| PAX8 | TGCCTCACAACTCCATCAGA | CAGGTCTACGATGCGCTG |
| ID2 | AGTCCCGTGAGGTCCGTTAG | AGTCGTTCATGTTGTATAGCAGG |
| CDX2 | GGGCTCTCTGAGAGGCAGGT | GGTGACGGTGGGGTTTAGCA |
| HHEX | CCTCTGTACCCCTTCCCG | GGGGCTCCAGAGTAGAGGTT |
| PDX1 | CGTCCGCTTGTTCTCCTC | CCTTTCCCATGGATGAAGTC |
| SCGB3A2 | GGCTAAGGAAGTGTGTAAATGAGC | CCATCCACCTCCGCTCTTTATC |
| TP63 | CCACAGTACACGAACCTGGG | CCGTTCTGAATCTGCTGGTCC |
| MUC5AC | GCACCAACGACAGGAAGGATGAG | CACGTTCCAGAGCCGGACAT |
| SCGB1A1 | ATGAAACTCGCTGTCACCCT | GTTTCGATGACACGCTGAAA |
| FOXJ1 | CAACTTCTGCTACTTCCGCC | CGAGGCACTTTGATGAAGC |
| SFTPB | TGCCTGGACCACCTCATCCTTG | GTCCTCACACTCTTGGCATAGG |
| AGER | GCCACTGGTGCTGAAGTGTA | TGGTCTCCTTTCCATTCCTG |
| PDPN | AGGAGAGCAACAACTCAACGGGAA | TTCTGCCAGGACCCAGAGC |
| SFTPC | AGCAAAGAGGTCCTGATGGA | CGATAAGAAGGCGTTTCAGG |
| LAMP3 | ACCGATGTCCAACTTCAAGC | TGACACCTTAGGCGGATTTT |
| CXCL8 | TTTTGCCAAGGAGTGCTAAAGA | AACCCTCTGCACCCAGTTTTC |
| CXCL10 | GTGGCATTCAAGGAGTACCTC | TGATGGCCTTCGATTCTGGATT |
| IL-6 | ACTCACCTCTTCAGAACGAATTG | CCATCTTTGGAAGGTTCAGGTTG |
| IL-1β | ATGATGGCTTATTACAGTGGCAA | GTCGGAGATTCGTAGCTGGA |
| TLR5 | GCCGGTCCTGTGTTTGGAAT | GGTGAGGTTGCAGAAACGATAAA |
| TNF-α | CCTCTCTCTAATCAGCCCTCTG | GAGGACCTGGGAGTAGATGAG |
| C3 | GGGGAGTCCCATGTACTCTATC | GGAAGTCGTGGACAGTAACAG |
| C2 | GGGGACAAGGTCCGCTATC | GAAGTCATAAGAGTAGGGTTGGC |
| CFB | GCACTGGAGTACGTGTGTCC | CCCGTTCTCGAAGTCGTGTG |
| CR2 | GGTCCTCGGGATTTCTTGTGG | GAACAACTGTACCTTATCACGGT |
| C4BPA | ATGACCTTGATCGCTGCTCTG | GTCAACGTAATATCCATCGGGG |

**Table S2.**

List of GO terms enriched in DEGs between FLA-PA-treated RAS cells and control RAS cells.

**Table S3.**

List of GO terms enriched in DEGs between RAS cells from COPD versus from healthy controls.

**Table S4.**

Cell type assignment. Overview of all gene signatures applied and generated in this study.

**Movie S1.**

Beating cilia were visible in apical-out d-AWO.
